# Tie2 signaling in the tumor microenvironment orchestrates breast cancer cell dissemination through TMEM doorways

**DOI:** 10.64898/2026.04.26.720938

**Authors:** Camille L Duran, Chinmay R. Surve, Prachiben P. Patel, Jack Hirsch, Jiufeng Li, Xianjun Ye, Nicole D. Barth, Xiaoming Chen, Suryansh Shukla, George S. Karagiannis, John C. McAuliffe, David Entenberg, Dianne Cox, John S. Condeelis, Maja H. Oktay

## Abstract

During breast cancer metastasis, tumor cells migrate toward intratumoral blood vessels and intravasate through stable structures known as TMEM (Tumor Microenvironment of Metastasis) doorways. TMEM doorways, composed of a Mena-expressing tumor cell, a Tie2^hi^/VEGF^hi^ macrophage, and an endothelial cell, are clinically validated prognostic markers of distant metastasis in breast cancer and represent the exclusive sites of tumor cell intravasation. We previously demonstrated that Tie2 signaling is essential for TMEM doorway function and tumor cell intravasation. In this study, we investigated how Tie2 signaling promotes tumor cell intravasation and metastasis. Because all three TMEM doorway-associated cell types can express Tie2, we sought to determine which of these cells contribute to the Tie2 signaling-dependent vascular opening at TMEM doorways and tumor cell dissemination. We found that endothelial cells associated with TMEM doorways secrete Ang2, which stimulates VEGF-A expression in Tie2^hi^ macrophages. Elevated VEGF-A levels at TMEM doorways increase vascular permeability, facilitating tumor cell entry into the bloodstream. Using tissue staining and line-scan analysis of Tie2 and lineage markers in human and mouse breast cancer models, we observed Tie2 expression in macrophages, tumor cells, and endothelial cells. To assess functional contributions, we selectively disrupted Tie2 in macrophages, endothelial cells, and cancer cells using CRISPR-Cas9 and RNAi approaches and tested in which of these cell-knockouts of Tie2 expression affected transendothelial migration *in vitro*. Macrophage-specific Tie2 deletion had the greatest impact on tumor cell intravasation. To confirm this finding *in vivo*, we generated a mouse model with inducible, macrophage-specific Tie2 knockout. Acute, targeted loss of Tie2 specifically in macrophages significantly reduced TMEM doorway associated vascular opening and tumor cell intravasation. Together, these findings establish macrophage Tie2 signaling as a critical driver of TMEM doorway-mediated vascular permeability and metastatic dissemination in breast cancer.

## Introduction

Breast cancer mortality is driven largely by distant metastasis, a process that requires tumor cells to access the vasculature, intravasate through discrete openings, and survive in the circulation as circulating tumor cells (CTCs). Tumor cell intravasation does not occur uniformly along the vessel wall but is instead exclusively restricted to specialized, microanatomical structures within the tumor microenvironment, termed Tumor Microenvironment of Metastasis (TMEM) doorways ^1, 2^. TMEM doorways are a tripartite cellular structure consisting of a Mena-expressing tumor cell, a perivascular VEGF-A expressing macrophage, and an endothelial cell, in direct and stable physical contact ^1, 3, 4^. TMEM doorway density has been clinically validated as a prognostic marker of distant metastatic recurrence in estrogen receptor-positive, HER2-negative breast cancer, independent of other clinical prognostics ^3–6^. Therefore, TMEM doorways represent both the physical location for tumor cell entry into the bloodstream and a promising potential therapeutic target for prevention of metastatic spread.

Tumor-associated macrophage (TAM) abundance and activation state strongly correlate with metastatic risk ^7–12^. TAMs are recruited to breast tumors through CSF-1/CSF-1R signaling and inhibition of this pathway delays macrophage recruitment, tumor progression, tumor cell dissemination and metastasis ^13–15^. Live cell and Intravital imaging have shown that TAMs promote tumor cell migration (streaming) toward TMEM doorways on blood vessels through a CSF-1/EGF signaling axis ^16–19^. In breast tumors, perivascular TMEM doorway macrophages are characterized by elevated VEGF-A expression, which when stimulated by CSF-1, promote local vascular permeability (TMEM doorway associated vascular opening, TAVO) and tumor cell intravasation through localized secretion of the accumulated VEGF-A ^1, 15^. However, a critical gap in information persists: the upstream signals that drive and maintain this elevated VEGF-A expression in TMEM doorway macrophages, and ensure its spatially restricted secretion at TMEM doorways, remain unknown.

A parallel body of work has identified a specialized subset of Tie2-expressing macrophages as potent regulators of tumor angiogenesis and vascular remodeling. Tie2 (also known as *TEK*) is a receptor tyrosine kinase expressed in endothelial cells, TAMs, and certain tumor cells, making it a critical regulator of the tumor microenvironment ^20^. During tumor progression, Tie2 signaling functions as an “angiogenic switch,” with tumor hypoxia driving the upregulation of both the Tie2 and its ligand, Angiopoietin-2 (Ang2) ^21–26^. Studies have shown that Tie2⁺ macrophages are enriched at areas of increased vascular density, “vascular hotspots,” where they respond to endothelial-derived Ang2 to support tumor revascularization following therapy and drive tumor growth and metastasis ^24, 27, 28^. In breast cancer and other malignancies, Ang2 levels are frequently elevated and pharmacologic Tie2 inhibition normalizes tumor vasculature and reduces metastatic dissemination across models ^24, 29, 30^.

Yet two fundamental questions remain unanswered: (1) which cell type, endothelial cell, macrophage, tumor cell, or some combination, mediates the pro-metastatic effects at sites of tumor cell intravasation, and (2) what are the precise cellular and spatial mechanisms by which Ang2/Tie2 signaling integrates with the CSF-1/VEGF-A axis that is known to control TMEM doorway opening and tumor cell dissemination. Despite the established role for Tie2 in angiogenesis and the clinical trials with Tie2 inhibitors, the spatial and mechanistic link between Ang2/Tie2 signaling and tumor cell intravasation and the Tie2 cell-type specific contributions to TMEM doorway opening are unknown. The work described here demonstrates the mechanism through which endothelial cell-derived Ang2 and macrophage Tie2 signaling in perivascular niches cooperate to control VEGF-A expression in TMEM doorway macrophages, TMEM doorway opening, and metastatic dissemination in breast cancer.

## Materials and Methods

### Cell culture

6DT1 murine breast cancer cells, RAW264.7 subline LR5 macrophages, and 3b-11 endothelial cells were cultured in Dulbecco’s Modified Eagle Medium (DMEM) (cat# SH30243, Hyclone, GE Healthcare Life Sciences, Logan, UT, USA) supplemented with 10% fetal bovine serum (FBS) (cat# S11550, Atlanta Biologicals, Flowery Branch, GA, USA). BAC1.2F5 macrophages^31^, bone marrow derived macrophages (BMMs), and immortalized BMMs (iBMMs) overexpressing Tie2 (Tie2^hi^ iBMMs) or control (Tie2^lo^ iBMMs) were cultured in Minimum Essential Medium, Alpha (α-MEM) (cat# 15-012-CV, Corning, Tewksbury, MA, USA) supplemented with 10-15% FBS (cat# 100-106, Gemini Bio-Products, Sacramento, CA, USA) and 36 µg/mL human recombinant CSF-1 (3000 units/mL, a gift from Dr. E. Richard Stanley). Generation of Tie2^hi^ and control (Tie2^lo^) iBMMs was previously described^30, 32^. Human Umbilical Vein Endothelial Cells (HUVECs) were cultured in complete EGM-2 (cat# CC-3162, Lonza, Allendale, NJ, USA) and were not used beyond passage five for any experiments.

### Cell line generation

Tie2 (*Tek*) was targeted using CRISPR-Cas9 technology in 3b-11 endothelial cells and 6DT1 breast cancer cells. *Tek*-targeting and non-targeting guide RNAs were cloned into the pLentiCRISPR-E (cat# 78852, Addgene, Watertown, MA, USA) and used to make lentiviruses using Lipofectamine 3000 (cat# L3000-008, Invitrogen). Cells were transduced with concentrated viral supernatants, single cell sorted, and selected with puromycin (0.5-3 μg/mL). Tie2 mRNA and protein knockdown were confirmed using qPCR and western blotting, respectively, comparing sgTie2 cells sgControl cells.

Tie2 was targeted using siRNA in RAW/LR5 macrophages (siTie2, siControl). RAW/LR5 cells were treated with AUM*silence* sdASOs (15 μM TEK-1, TEK-2, or non-targeting) targeting Tie2 or control sequence for 72 hours (AUM Biotech, Philadelphia, PA, USA). qPCR and western blotting were used to confirm Tie2 mRNA and protein knockdown, respectively.

Bone marrow-derived macrophages (BMMs) were isolated from the tibia and femurs ^33^ of Tie2^+/+^, Tie2^fl/fl^/*Csf1r-Mer-iCre-Mer^(-)^* (-Cre), or Tie2^fl/fl^/*Csf1r-Mer-iCre-Mer^(+)^* (+Cre) mice that had been treated with or without corn oil or 4-OH tamoxifen (described in “*in vivo* drug administration studies” section, below). Tie2^+/+^/-Cre, Tie2^fl/fl^/-Cre, or Tie2^fl/fl^/ +Cre BMMs were grown in alpha-MEM supplemented with 15% FBS, 36 ng/mL CSF-1, and antibiotics. Tie2 knockdown in Tie2^fl/fl^/ +Cre BMMs was confirmed using western blotting.

Generation of immortalized BMMs overexpressing Tie2 (BMM-Tie2^hi^) and control (BMM-Tie2^lo^) has been previously described^30, 32^.

### Mice

All studies involving mice were conducted in accordance with the National Institutes of Health regulations concerning the care and use of experimental animals. The procedures were approved by the Albert Einstein College of Medicine Institute for Animal Care and Use Committee. FVB/N-Tg(MMTV-PyVT)634Mul/J mice (MMTV-PyMT, stain #002374) and *SCID* mice (strain #001303) were purchased from The Jackson Laboratory (Bar Harbor, ME, USA). Transgenic mice expressing the PyMT antigen and dendra2 fluorescent protein under the MMTV promoter, MMTV-iCre/CAGCAT-Dendra2/MMTV-PyMT ^34^ were bred in house.

Tie2-floxed (Tie2^fl^) mice were generated with the help of the Transgenic Mouse Core at Albert Einstein College of Medicine. Two CRISPR guide RNAs were designed to mediate insertion of loxP sites flanking exon 1 of Tie2 (*Tek*), gRNA 64-41; GAAAACTTTAAGCTTGGTAT TGG and gRNA IN1 72-65; CTCTCTAGAGGTGCCACTAC AGG. Cas9 protein, the two gRNAs, and single-stranded homology-directed repair (HDR) donors (Tie2 5′ HRD and Tie2 3′ HRD), each containing a loxP site flanked by ∼40-nt homology arms, were co-injected into fertilized FVB eggs. Homology-directed repair at each cut site introduced a 5′ loxP immediately upstream of exon 1 and a 3′ loxP within intron 1, generating a floxed exon 1. Tie2^fl/+^ mice were crossed with wild-type FVB to obtain F1 Tie2^fl/+^ heterozygotes and subsequently intercrossed to generate Tie2^fl/fl^ homozygous mice for experimental use. Genotyping on Tie2^fl/fl^ litters was performed using primers: 5’ ccagttaagggcttttcgtc 3’ and 5’ aaccaattcggggaatccta 3’, and 5’ ggataacaaactctgggagca 3’ and 5’ tgaggccctgtctcaaaact 3’. To create macrophage-specific Tie2-inducible knockout mice, Tie2^fl/fl^ mice were crossed with the *Csf1r-Mer-iCre-Mer ^(+)^* mice ^35^, bred in house.

### in vivo drug administration studies

Inhibitors were administered based on established doses of efficacy found in previous studies. In rebastinib inhibitor experiments, tumor pieces from *MMTV-PyMT/dendra2* mice or HT17 human PDX tumor chunks were orthotopically transplanted into FVB or *SCID* mice, respectively, as previously described ^36–38^. Following about four weeks of tumor growth, animals were divided into two groups. Animals were treated with the Tie2 inhibitor, rebastinib, 10 mg/kg *p.o.* (100 µL total volume) or HPMC (Vehicle, 100 µL total volume) twice per week, for four weeks. Rebastinib was kindly provided by Deciphera Pharmaceuticals and was reconstituted at a concentration of 10 mg/mL in 0.4% hydroxypropyl methylcellulose (HPMC).

In experiments to induce Cre recombinase in Tie2^fl/fl^ animals, tumor chunks from MMTV-PyMT animals were orthotopically transplanted into Tie2^fl/fl^/*Csf1r-Mer-iCre-Mer^(-)^* (-Cre) or Tie2^fl/fl^/*Csf1r-Mer-iCre-Mer^(+)^* (+Cre) and once the tumors reached at least 0.6 cm^3^, animals were further divided into two more groups. Tie2^fl/fl^/-Cre or Tie2^fl/fl^/+Cre mice were treated at 96 and 72 hours before sacrificing with either 50 mg/kg 4-OH tamoxifen (+Tam, 100 µL total volume) or 100 µL of an ethanol-corn oil solution (-Tam, 10% Ethanol, 90% corn oil). 4-OH tamoxifen (Sigma, H6278) was reconstituted first in ethanol and then in corn oil, with a final concentration of 10 mg/mL in 10% ethanol, 90% corn oil. One cohort of experimental mice was used for flow experiments, detailed below. A second cohort was used for circulating tumor cell (CTC) and primary tumor staining experiments. In this second cohort, mice were injected *i.v.* with 20 mg/mL 155 kDa TMR-dextran (cat# T1287, Sigma-Aldrich, Burlington, MA, USA) diluted in PBS, 15 minutes before the termination of the experiments. To collect circulating tumor cells (CTCs), mice were anesthetized with isoflurane, and blood was collected from the right ventricle via cardiac puncture with a heparinized syringe and incubated with RBC lysis buffer (cat# 00-4333-57, Invitrogen) for five minutes and neutralized with PBS. Cells were spun at 300xg for 10 minutes at 4°C and pellets were suspended and cultured in DMEM/F12 (cat# 11320033, Gibco) supplemented with 20% FBS. Adherent tumor cells were quantified as CTCs at time of no tumor cell growth, as previously described ^1, 39^. The raw number of CTCs for each mouse was normalized to the volume of blood collected for each mouse. The number of CTCs in the +Tam treatment group was then set relative to the average number of CTCs in the -Tam control group. Primary breast tumors were collected at the time of sacrifice and fixed in either 10% formalin or 1% paraformaldehyde and used for subsequent tissue staining.

### in vitro VEGF-A expression studies

For *in vitro* Tie2 inhibition studies, rebastinib was dissolved in DMSO and used at concentration of 50 nM. In macrophage co-culture experiments, 1x10^5^ BAC1.2F5 macrophages, labeled with CellTracker Green, were serum starved overnight in basal medium supplemented with 0.5% FBS and 300 units/mL of CSF-1 (one-tenth the amount of CSF-1 added to complete culture media). BAC1.2F5 macrophages were then cultured alone, with HUVECs, or with media conditioned (CM) by HUVECs for 24 hours. Cultures were treated with DMSO control or 50 nM rebastinib for 24 hours. In macrophage Ang2 stimulation experiments, serum starved BAC1.2F5 macrophages, as described above, were pretreated with 50 nM rebastinib or DMSO. BMMs treated with siRNA against Tie2 or control (see transfection description above) and serum starved overnight in basal medium supplemented with 0.5% FBS. Macrophages were then cultured overnight with 250 ng/mL Ang2. In macrophage Ang2 and CSF-1 stimulation experiments, BAC1.2F5 macrophages were serum-starved, as described above, and then pretreated with 50 nM rebastinib or DMSO. Macrophages were then cultured overnight with 250 ng/mL Ang2 (cat# 7186-AN-025, R&D Systems) and the next day, macrophages were treated with or without 3000U/mL CSF-1 for 45 minutes. In all experiments above, at the completion of experiments, cells were washed in cold PBS, fixed in 4% paraformaldehyde for 20 minutes, permeabilized in 0.1% Triton-X 100, and blocked. Cells were stained with antibodies against VEGF-A (cat# 512809, clone 2G11-2A05, Biolegend) and Alexa Fluor647-conjugated secondary antibodies, and imaged. In co-culture experiments, macrophage VEGF-A expression was measured by outlining the CellTracker Green labeled cells and only quantifying the VEGF-A signal (far red, 647nm channel) contained within those macrophage outlines, using the FIJI (NIH) pixel intensity measurement. In all mono-cell culture experiments, VEGF-A macrophage expression was measured using the FIJI (NIH) pixel intensity measurement.

### Tissue harvesting and immunofluorescence labeling of vascular opening (TMEM doorway-associated vascular opening) with 155 kDa dextran-TMR

*In vivo* labeling of the flowing vasculature and sites of vascular opening was performed as previously described ^1^. Briefly, to quantify vascular opening events (TMEM doorway associated vascular opening), high molecular weight 155 kDa TMR-dextran (cat# T1287, Sigma-Aldrich, Burlington, MA, USA) diluted in PBS to 20 mg/mL was administered by tail vein *i.v.* fifteen minutes before termination of the experiments. At time of sacrifice, primary tumors and lungs were harvested and split to be used for frozen sections or paraffin-embedded sections. Tissues to be used for frozen sections were fixed in 1% paraformaldehyde for 24 hours at 4°C, then added to a 30% sucrose solution at 4°C, and finally embedded in OCT. Tissues used for paraffin-embedded sections were fixed for 48 hours in 10% formalin in a volume ratio for tumor to formalin of 1:20 and then embedded in paraffin blocks. Frozen sections and paraffin blocks of tumors were cut into 10 µm sections and immunofluorescence staining was performed. TMR-Dextran is visualized using rabbit anti-TMR (A-6397; Life Technologies, Carlsbad, CA, USA).

### Immunofluorescence staining of tissue

Tumor sections were dewaxed and rehydrated in alcohol followed by water. Antigen retrieval was performed with a citrate solution at pH 5.5 or EDTA pH 9.0, depending on the primary antibody, in a humidified chamber. For TSA/OPAL staining, endogenous HRP was blocked after a ten-minute incubation with 30% hydrogen peroxide. Slides were washed with PBS-T (0.1% or 0.05% Tween-20) and blocked (1% BSA, 10% FBS, 1% goat serum in PBS-T) for 1 hour at room temperature. TMEM doorways were stained using IHC with three antibodies: anti-Mena (cat# NBP1-87914, Novus Biologicals); anti-endomucin (cat# sc-65495, Santa Cruz Biotechnology, Dallas, TX, USA); and anti-Iba1 (cat# 019-19741, FUJIFILM Wako Chemicals, Richmond, VA), applied sequentially and developed separately with different chromogens on a Dako Autostainer ^4, 5, 30^. The sequential tissue sections were stained using immunofluorescence with different combinations of: anti-endomucin (cat# sc-65495, Santa Cruz Biotechnology), anti-TMR (to visualize TMR-dextran, cat# A-6397, Thermo Fisher Scientific), anti-ZO-1 (cat# 402200, Invitrogen), anti-CD31 (cat# 77699; Cell Signaling Technology, Danvers, MA, USA), anti-Iba1 (cat# 019-19741, FUJIFILM Wako Chemicals), anti-VEGF-A (cat#512809, clone 2G11-2A05, Biolegend, San Diego, CA, USA) or anti-VEGF-A (cat# MA5-32038,Thermo Fisher), Alexa Fluor 546-conjugated anti-Ang2 (cat# sc-74403 AF546, Santa Cruz Biotechnology), anti-Tie2 (cat#14598782, Ebioscience, San Diego, CA, USA), anti-Ly-6G (cat# 87048, Cell Signaling Technology), anti-CD68 (cat# MCA1957, clone FA-11, Serotec), and anti-F4/80 (cat# 70076, Cell Signaling Technology). Sections were washed with PBS-T and the primary antibodies were detected with Alexa Fluor 488, 555 or 647 conjugated secondary antibodies targeting the primary antibody species (Invitrogen, Eugene, OR, USA) or detected with HRP-conjugated secondary antibodies and Opal-TSA fluorophores (cat# NEL810001KT, Akoya Biosciences, Malborough, MA, USA). Nuclei were stained with 4, 6-diamidino-2-phenylindole (DAPI). Fluorescently labeled samples were mounted with Prolong Gold or Prolong Diamond antifade reagent (cat# P36930, P36961, Invitrogen) and imaged with a PerkinElmer Pannoramic 250 Flash II digital whole-slide scanner using a 20x 0.8NA Plan-Apochromat objective (PerkinElmer, Hopkinton, MA, USA). Images of individual fields of view or whole slides were imported into ImageJ or VisioPharm for analysis.

### Extravascular dextran, Ang2, and VEGF-A quantification at TMEM doorways

We used a previously described method to identify active TMEM doorways within the entire tissue section that takes into account the presence of high-molecular weight dextran that extravasates into the tissue, as a result of TMEM doorway associated vascular opening (TAVO) ^15, 40^. In short, serial tissue sections were cut and stained for TMEM doorways by staining for Iba1, endomucin, and Mena by IHC (tissue section 1), and for TMR-Dextran, Ang2 or VEGF-A, Iba1 or F4/80 to identify macrophages, and endomucin by IF (tissue section 2). F4/80 and Iba1 antibodies are used to identify macrophages and have been found to be equivalent and interchangeable. The sections were then imaged with a digital whole slide scanner and aligned to the single cell level using the TissueAlign module in Vis (Visiopharm, Hoersholm, Denmark). IF images were thresholded above background, creating binary masks for endomucin (blood vessel) and dextran signals. These masks were then superimposed to differentiate between intravascular (overlapping with the endomucin mask) and extravascular (outside the endomucin mask) dextran signal. In the IHC section, TMEM doorways were identified using previously published criteria ^4, 5^ and superimposed onto the aligned IF stained slide, enabling the quantification of the amount of extravascular dextran within TMEM doorways. TMEM doorways were then defined as inactive or active TMEM doorways based on established thresholds of extravascular dextran contained within each doorway ^40^. To measure the intensity of extravascular Ang2 within TMEM doorways, Ang2 signal within the extravascular dextran mask (described above) was quantified within the inactive and active TMEM doorway ROIs in the IF slide. For quantification of intracellular VEGF-A and VEGF-A expression within TMEM doorways, the TMEM doorway associated macrophage was identified using the F4/80 stain in IF section, within the TMEM doorway circle, and an ROI mask was created. VEGF-A expression within the TMEM doorway macrophage ROI and outside the TMEM doorway macrophage ROI, but within the TMEM doorway circle, was then quantified.

### Tie2+ macrophage distance analysis relative to TMEM doorways and blood vessels without TMEM doorways, in fixed tissue in vivo

We used a previously described method to measure the distance of Tie2-expressing macrophages from the nearest TMEM doorway and nearest blood vessel without a TMEM doorway ^41^ In short, serial tissue sections were cut and stained for TMEM doorways by staining for Iba1, endomucin, and Mena by IHC (tissue section 1), and for Iba1 to identify macrophages, endomucin, and Tie2 by IF (tissue section 2). The sections were then imaged with a digital whole slide scanner and aligned to the single cell level using the TissueAlign module in Vis (Visiopharm, Hoersholm, Denmark). IF images were thresholded above background, creating binary masks for Iba1 (macrophages), Tie2, and endomucin (blood vessel) signals to identify Tie2 expressing macrophages. In the IHC section, TMEM doorways were identified using previously published criteria ^4, 5^ and superimposed onto the aligned IF stained slide. Each TMEM doorway was visualized as a 60 μm diameter circle and the distance of all Tie2+ macrophages per field were measured to its nearest TMEM doorway circle. As previously described, Tie2+ macrophages that were inside or touching the TMEM doorway boundary were counted in data point 0 μm in Figure 4E and 4I ^41^. Distance analysis of each Tie2+ macrophage to the nearest blood vessel without a TMEM doorway was performed by thresholding the endomucin stained blood vessel channel and removing blood vessels that contain a TMEM doorway. Tie2+ macrophages touching the blood vessel were counted in data point 0 μm in Figure 4F.

### Quantitative real-time polymerase chain reaction (qPCR)

To quantify gene expression in Tie2hi BMMs transfected with Tie2 siRNA or control siRNA and 6DT1 tumor cells and 3b-11 endothelial cells transduced with CRISPR-Cas9 Tie2-targeting lentiviruses, cells were grown under normal culture conditions. Total RNA was isolated from cells by using the RNeasy Plus Mini Kit (cat# 74134, Qiagen, Germantown, MD, USA) and cDNA was synthesized and amplified from 1 μg total RNA using the using iScript cDNA synthesis kit (cat# 1708891, BioRad) per manufacturer’s protocol. The qPCR was performed with SYBR Green PCR Master Mix (cat# 4367659, Thermo Fisher Scientific) using a QuantStudio 3 real-time PCR instrument (applied biosystems, Thermo Fisher Scientific). The following primers were used: mouse GAPDH 5’- CTCATGACCACAGTCCATGC -3’, 5’- CACATTGGGGGTAGGAACAC -3’; mouse VEGF-A 5’- AGCAGAAGTCCCATGAAGTGA -3’, 5’- ATGTCCACCAGGGTCTCAAT -3’; mouse Tie2 5’- TTCATCCACTCAGTGCCCCG -3’, 5’- CTGAGCTTCACATCTCCGAACA -3’. The mean cycle threshold (Ct) values were then used to analyze relative expression. Analysis was performed using comparative-CT method (2^−ΔCT^ method) and all Ct values were normalized to GAPDH. Each reaction was performed in triplicate.

### ELISA

In macrophage simulation experiments were set up as described in the “*in vitro* VEGF-A expression studies” section, above, and the figure legends. Media conditioned by the cultured cells were collected at the indicated time points and stored at -80°C until use. ELISAs were performed per the manufacturer’s recommendations using the Mouse VEGF-A DuoSet ELISA kit (cat# DY493, R&D Systems). The concentration of protein secreted was interpolated from the standard curve measurements.

### Trans-endothelial Migration Assay (iTEM assay)

The iTEM assay was performed as previously described ^15, 42^. The transwell (cat# 353097; 8 µm pore size, Corning, Corning, NY, USA) was prepared so that tumor cell trans-endothelial migration was in the intravasation direction found *in vivo* (from subluminal/tissue side to luminal side of the endothelium). To prepare the endothelial monolayer, the underside of each transwell was coated with 70 µL of Matrigel (2.5 µg/mL; cat# 354230, Corning). Approximately 5x10^4^ HUVECs or 3b-11 cells were plated on the Matrigel coated underside of the transwell. Transwells were then flipped into a 24-well plate containing 200 µL of compete EGM-2 (HUVECs) or DMEM (3b-11 cells) and monolayers were formed over a 24-hour period. The integrity of the endothelium was measured using low molecular weight dextran as described previously ^43^, and only confluent, non-permeable monolayers were used for the following steps. Macrophages and tumor cells were separately labelled with cell tracker dyes (cat# C7025, C34552, Invitrogen) before the experiment. Then, 1.5x10^4^ tumor cells (serum starved overnight in DMEM supplemented with 0.5% FBS) and 5x10^4^ macrophages were added to the upper chamber of the transwell in 200 µL of DMEM without serum while the bottom chamber contained EGM-2 (HUVECs) or DMEM (3b-11 cells) supplemented with 36 µg/mL of CSF-1. In rebastinib pretreatment experiments, 3b-11 endothelial cells, 6DT1 tumor cells, and RAW/LR5 macrophages were pretreated separately, overnight, with 50 nM rebastinib or DMSO control. The next morning, cells were rinsed three times with DPBS and then added to the assay.

After 18 hours, the transwells were fixed and stained for ZO-1 (cat# 402200, Invitrogen). Transwells were imaged using a Leica SP5 or SP8 confocal microscope using a 20x 0.75 NA objective and processed using ImageJ/Fiji ^44^. Tumor cell trans-endothelial migration quantification was performed by counting the number of tumor cells that had crossed the intact endothelium (intact monolayers confirmed by ZO-1 staining for tight junctions) within the same field of view (20x, three-four random fields per well, three-four wells per experiment) and represented as normalized values from at least three independent experiments.

### Flow cytometry experiments

Tie2 expression in cells of the tumor microenvironment was characterized by flow cytometry. Tumors from Tie2^fl/fl^/*Csf1r-Mer-iCre-Mer^(-)^* or Tie2^fl/fl^/*Csf1r-Mer-iCre-Mer^(+)^* mice treated with corn oil or 4-OH tamoxifen were harvested, rinsed in cold PBS, minced, and digested in lysis buffer containing DMEM, 100 μg/mL liberase TM (cat# 5401119001, Roche, Indianapolis, IN, USA), and 100 μg/mL DNAase I (cat# 90083, Invitrogen) for 25 minutes, rotating at 37°C. Lysates were rinsed in FACS buffer (containing PBS with 2% FBS) and filtered through a 40 μm strainer (cat# 352340, Corning). Lysates were spun down and incubated in RBC lysis buffer for 5 minutes on ice. Cells were rinsed three times in FACS buffer and stained with Live/Dead viability marker, Zombie UV (1:500 dilutionin PBS, cat# 423108, BioLegend) for 20 minutes on ice in the dark. Cells were then washed and resuspended in FACS buffer and Fc blocked with CD16/CD32 antibodies (cat# 14-0161-82, Invitrogen) for 15 minutes. Cells were then incubated for 45 minutes with antibodies directed against CD11c (cat# 117353, BioLegend), CD64 (cat# 139309, BioLegend), CD31 (cat# 160205, BioLegend), CD11b (cat# 101217, BioLegend), Ly-6G (cat# 127618, BioLegend), F4/80 (cat# 123112, BioLegend), CD115 (cat# 135528, BioLegend), CD202b/Tie2 (cat# ab95722, Abcam), Ep-CAM1 (cat# 118218, BioLegend), Ly-6C (cat# 566987, BD Biosciences), and CD45 (cat# 1115660, Sony Biotechnology, San Jose, CA, USA). Cells were washed three times with FACS buffer, resuspended in FACS buffer and filtered through a 40 μm cell strainer (cat# 352235, Corning). Samples were run on a Cytek Aurora spectral flow cytometer equipped with five lasers and compensation was applied using single color control OneComp eBeads compensation beads (cat# 01-1111-42, Invitrogen). FlowJo version 10.10.0 (BD Biosciences) was used for data analysis with gating strategies guided by IgG (cat# 400557 (BioLegend), 12-4301-82 (Thermo Fisher), 400523 BioLegend)), and fluorescence minus one (FMO) controls. Gating strategies are demonstrated in Supplemental Figure 5.

### Statistical analysis

Individual animals in each cohort are presented as individual points on a dot plot. A horizontal line indicates the mean value and the error bars represent the standard deviation or standard error of the mean, as indicated in the figure legend. Statistical significance was determined using an unpaired Student’s *t*-test, one-way ANOVA, or two-way ANOVA, with Tukey’s multiple comparisons test, as indicated, using GraphPad Prism (version 10; Graph Pad Software, La Jolla, CA). Data sets were checked for normality (D’Agostino & Pearson omnibus normality test or Shapiro-Wilk normality test) and unequal variance using GraphPad Prism. Welch’s correction was applied to *t*-tests as needed. *P* values less than 0.05 were deemed significant. For *in vitro* experiments, results are representative of at least three independent experiments.

## Results

### Extravascular Ang2 is increased at active TMEM doorways

Ang2 is produced by endothelial cells and can signal through Tie2 to regulate tumor cell dissemination and metastasis^24, 27, 28^. We hypothesized that Ang2 released from endothelial cells at TMEM doorways promotes vascular opening and thereby contributes to TMEM doorway function. To test this, we first asked whether Ang2 levels are elevated in TMEM doorway-associated endothelial cells at *active* TMEM doorways, defined as TMEM doorways associated with a vascular opening that permits extravasation of high molecular weight dextran into the perivascular space **(Figure 1A)**.

**Figure 1:**
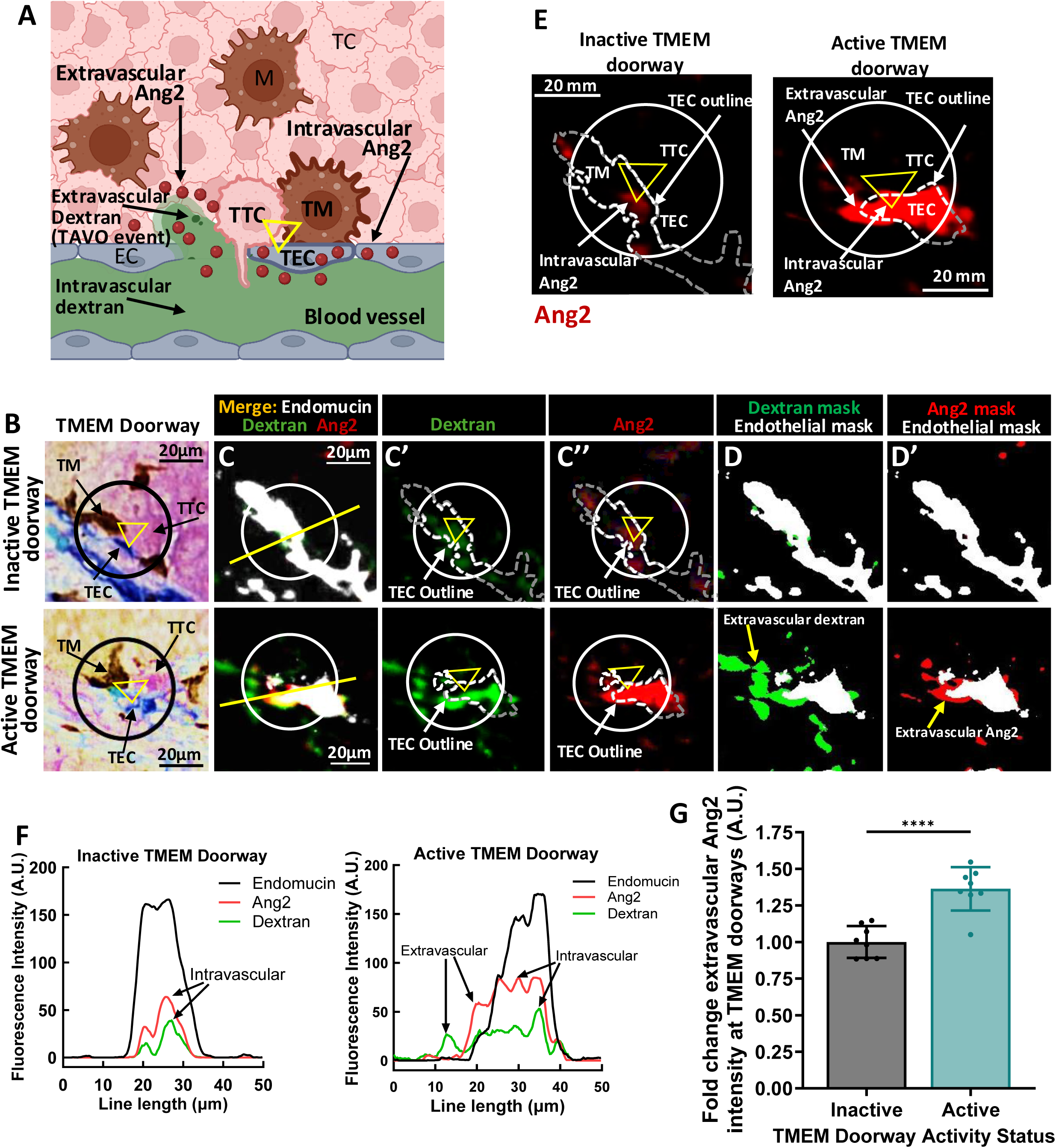
Extravascular Ang2 is increased at active TMEM doorways. **A)** Cartoon denoting the identification of parameters used for analysis of Ang2 levels in TMEM doorways and TMEM doorway activity. The three cells composing the TMEM doorway are indicated by the yellow triangle connecting the TMEM doorway macrophage (TM), the TMEM doorway endothelial cell (TEC) and the TMEM doorway tumor cell (TTC). Intravascular dextran (green) and intravascular Ang2 (red circles) is in the lumen of blood vessel and TECs, extravascular dextran and extravascular Ang2 is outside the lumen of blood vessel. Figure created with BioRender.com. **B)** Panel shows inactive and active TMEM doorways, visualized by immunohistochemistry (IHC) staining for Mena, Iba-1, and endomucin. The three cells of the TMEM doorway (contained in black circle, with the three cells forming the TMEM doorway indicated with the yellow triangle) are the TMEM doorway endothelial cell (TEC, endomucin stained in blue), TMEM doorway macrophage (TM, Iba1 stained in brown), and TMEM doorway tumor cell (TTC, Mena stained in pink) and black arrows indicate where the cells are localized within the TMEM doorway. TMEM doorways were identified using automated analysis by VisioPharm identifying three adjacent immuno-histochemical stains. **C)** The sequential tissue sections after the IHC in (B) were stained using immunofluorescence (IF) with antibodies against endomucin (white), dextran (green), Ang2 (red). The two sequential sections from 1B and 1C were aligned and the same TMEM doorways were matched between the two sections, as indicated by the black circle in the IHC panel (1B) and white circle in the IF panels (1C). **(c’)** Shows a white dotted outline of the TECs and the green dextran signal outside the TEC outline is the extravascular dextran. **(c’’)** The white TEC outline is overlaid on the red Ang2 signal, showing the extravascular Ang2 outside of the outline. The panels demonstrate the association of Ang2 level in TMEM doorways with TMEM doorway activity. **D)** Shows the extravascular signal for dextran as a green mask and the endomucin stain (TEC) as a white mask, where thresholded, positive signal for these stains was converted into a binary mask. Active versus inactive TMEM doorways were distinguished by the presence of extravascular dextran staining (non-overlapping with the endomucin stain), which indicates that the vessel had a TMEM doorway-associated vascular opening (TAVO). **d’)** Shows endomucin mask (white) and the extravascular Ang2 stain converted into a binary mask (red), demonstrating the presence of extravascular Ang2 in active TMEM doorways. Scale bars=20 μm. **E)** High magnification images from panel c”, showing the position of Ang2 (red) relative to the TMEM doorway endothelial cell (TEC) (dashed white outlines) in inactive (upper) and active (lower) TMEM doorways. Scale bars 20μm. **F)** Quantification of the immunofluorescence intensity of Ang2, dextran and endomucin across the yellow line scan across the TMEM doorway from Merge IF stain in (C). **G)** Immunofluorescence measurement of Ang2 levels in active and inactive TMEM doorways. TMEM doorways were identified using the IHC stained TMEM doorway slide as in (1B and 1C) and the active and inactive TMEM doorways were identified in the sequential IF-stained sections as described above (see panel C). The level of extravascular Ang2 in the TMEM doorways was measured in both active and inactive TMEM doorways using Visiopharm. Between 82-134 TMEM doorways were analyzed per mouse, in 8 mice. Each point represents the average fold change Ang2 intensity in inactive or active TMEM doorways in one mouse, bars show the fold change mean intensity (A.U.) and error bars represent standard deviation. \*\*\*\**P* < 0.0001 analyzed by Student’s *t*-test.

We performed multiplex staining of breast tumor sections from polyoma middle T antigen (PyMT) mice injected intravenously with high-molecular weight dextran (155 kDa, green) 1 hour before sacrifice. We stained two sequential tumor sections and digitally aligned them at single-cell resolution: one section was stained by IHC for TMEM doorway components (Iba1, Endomucin, Mena) to identify TMEM doorways **(Figure 1B, IHC stained panel)**, and the adjacent section was stained by IF for Ang2, dextran, and endothelial cells (Endomucin) **(Figure 1C, IF stained panels)**. TMEM doorways were then classified as inactive (upper panels) or active (lower panels) based on the presence of dextran in the extravascular space adjacent to the TMEM doorway, as previously described ^1, 37^.

At inactive TMEM doorways, Ang2 signal was confined to the intravascular compartment of endothelial cells, whereas at active TMEM doorways Ang2 was detected both intra- and extravascularly **(Figure 1D-E, IF stained panels)**, consistent with Ang2 secretion from endothelial cells at active TMEM doorways. Using line profiles drawn across inactive and active TMEM doorways (yellow lines to define areas with extravascular dextran, **Figure 1C**), we observed that extravascular dextran signal co-localized with elevated extravascular Ang2 at active TMEM doorways **(Figure 1F)**. Quantification of extravascular Ang2 (red) showed that its fluorescence intensity was higher in active compared with inactive TMEM doorways **(Figure 1G)**, supporting a role for Ang2 in regulating TMEM doorway associated vascular opening.

### Endothelial cells induce an increase in intracellular VEGF-A in macrophages via Tie2/Ang2 paracrine signaling

We previously reported that macrophage-derived VEGF-A is a critical regulator of TMEM doorway opening ^1^ and its secretion is triggered by tumor cell-derived CSF-1 signaling through the CSF-1R on macrophages ^15^. However, CSF-1/R signaling did not account for the elevated expression of VEGF-A^15^ in macrophages of the TMEM doorway. Because perivascular macrophages can also express the Tie2 receptor ^1, 24, 27, 28^, we tested whether endothelial cell-secreted Ang2 acting on macrophage Tie2 regulates macrophage VEGF-A expression. To distinguish between intracellular and secreted VEGF-A, we used VEGF-A immunofluorescence (IF) to assess intracellular levels of VEGF-A, and a VEGF-A ELISA on conditioned media to measure its secretion.

We first quantified intracellular VEGF-A in BAC1.2F5 macrophages co-cultured with human umbilical vein endothelial cells (HUVECs), labeling macrophages with CellTracker Green to distinguish them from endothelial cells **(Figure 2A)**. In endothelial cell–macrophage co-cultures, intracellular macrophage VEGF-A levels were significantly higher than in macrophages cultured alone (**Figure 2B**, middle panel). To determine whether this effect was mediated by paracrine rather than juxtacrine signaling, we cultured macrophages in HUVEC-conditioned media (CM) and observed a similar increase in intracellular macrophage VEGF-A, consistent with a paracrine mechanism (**Figure 2B**, bottom panel).

**Figure 2:**
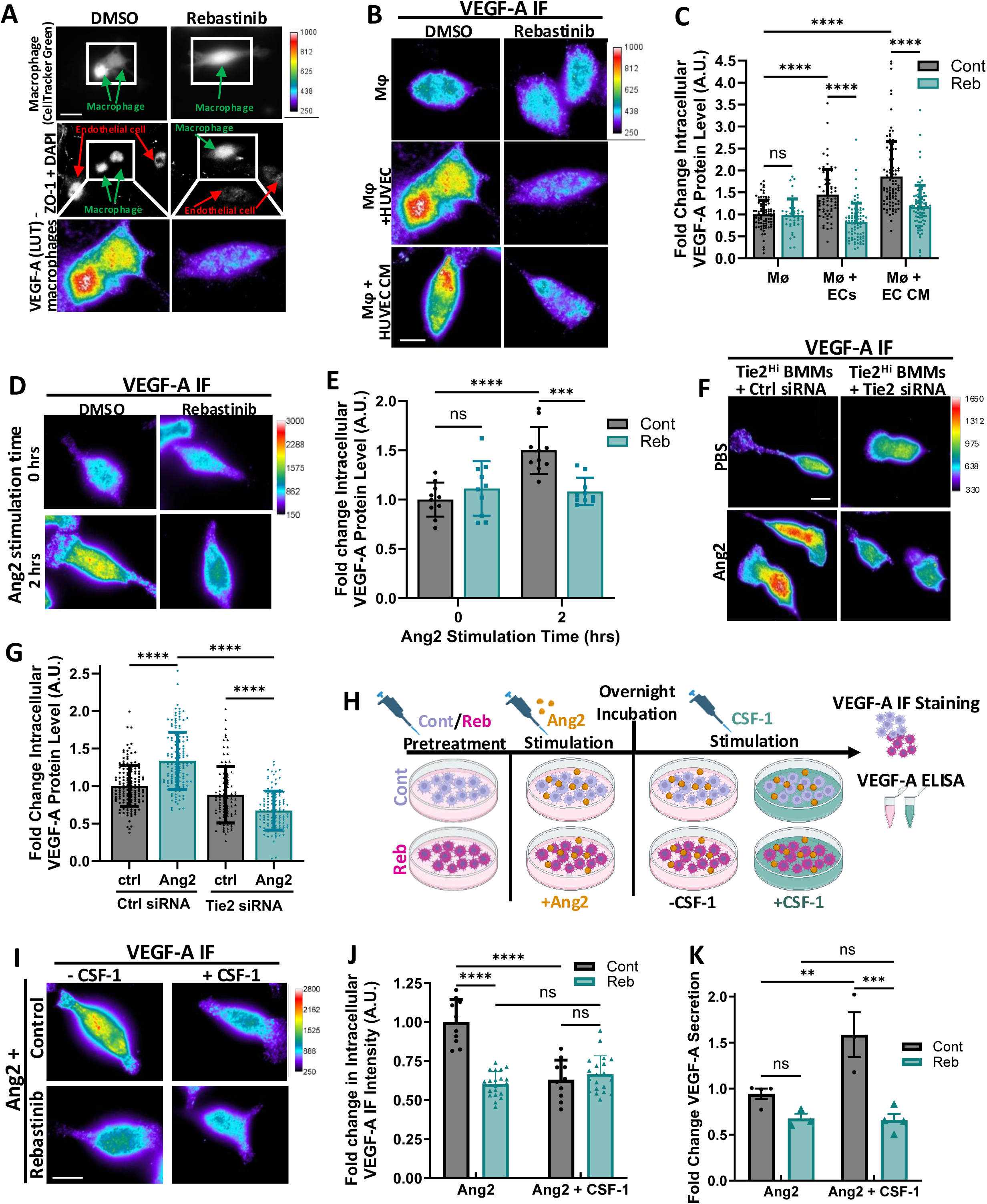
Endothelial cells induce an increase in intracellular VEGF-A in macrophages via Ang2-Tie2 paracrine signaling. **A)** Endothelial cells and macrophages were co-cultured to measure induction of VEGF-A protein expression. Macrophages were labelled with CellTracker^TM^ Green to distinguish them from endothelial cells (top panel), the middle panel shows ZO-1 and DAPI stain to identify the endothelial cells and nuclei of both cell types respectively, and the bottom panel is magnified from the boxed region shown in middle panels, showing VEGF-A staining in macrophages in pseudo color LUT. Scale bars 10 µm. **B)** Immunofluorescence staining of VEGF-A (pseudo color) in macrophages (BAC1.2F5, M⏀) co-cultured with endothelial cells (HUVECs) or endothelial cell conditioned media (CM) in the presence of DMSO or Tie2 inhibitor rebastinib (50nM), as described in (A). Scale bars 5 µm. **C)** Quantification of immunofluorescence intensity of intracellular VEGF-A in macrophages from (B) n=3 experiments, \*\*\*\**P* < 0.0001 analyzed by two-way ANOVA showing the inhibition of macrophage VEGF-A expression by Rebastinib. **D)** Immunofluorescence staining of VEGF-A (pseudo color) in macrophages (BAC1.2F5) stimulated with Ang2 (250ng/ml) in presence of DMSO or Tie2 inhibitor rebastinib (50nM). Scale bars 5 µm. **E)** Quantification of immunofluorescence intensity of intracellular VEGF-A in macrophages from (D) showing Tie2 signaling induces VEGF-A expression in macrophages and inhibition of Tie2 by Rebastinib decreases VEGF-A expression. n=3, \*\*\**P* < 0.001, \*\*\*\**P* < 0.0001 analyzed by two-way ANOVA. **F)** Immunofluorescence staining of VEGF-A (pseudo color) in macrophages (BMMs) transfected with ctrl siRNA or Tie2 siRNA stimulated with PBS or Ang2 (250ng/ml) as in (D). Scale bars 5 µm. **G)** Quantification of immunofluorescence intensity of intracellular VEGF-A in macrophages from (F) showing Tie2 knockdown in macrophages inhibits VEGF-A expression. n=3, \*\*\*\**P* < 0.0001 analyzed by two-way ANOVA. **H)** Experimental diagram of macrophage treatment for data shown in (I-K). BAC1.2F5 macrophages were pretreated with DMSO control (Cont) or 50nM Rebastinib (Reb) and then cultured overnight with 250ng/ml Ang2. The next day, macrophages were treated with or without 3000U/mL CSF-1 for 45 minutes, fixed, stained for VEGF-A (I-J) and conditioned media was used for VEGF-A ELISA (K). **I)** VEGF immunofluorescence staining of BAC1.2F5 macrophages from (H), Scale bar 5 µm. **J)** Quantification of immunofluorescence intensity of intracellular VEGF-A in macrophages (I) showing VEGF-A expression within macrophages decreases with CSF-1 treatment. n=3, ****p<0.0001 analyzed by two-way ANOVA. **K)** VEGF-A ELISA from conditioned media obtained from macrophages from (H), denoted as fold change of secreted protein compared to control showing that CSF-1 signaling increases VEGF-A secretion and this is blocked with Tie2 inhibition n=3, **p*<* 0.001, ***p<0.001, ns=not significant, analyzed by two-way ANOVA.

To test whether this increase in intracellular VEGF-A was driven by Ang2/Tie2 signaling, we inhibited Tie2 using rebastinib, a highly selective Tie2 inhibitor ^30^. Rebastinib abolished the increase in intracellular macrophage VEGF-A in both co-culture and HUVEC CM conditions, indicating that the VEGF-A upregulation in macrophages depends on Ang2/Tie2 signaling **(Figure 2B**, right panels; **Figure 2C)**. Consistently, direct stimulation of macrophages with recombinant Ang2 increased intracellular VEGF-A, and this effect was blocked by rebastinib **(Figure 2D, E)**. To further determine the involvement of macrophage Tie2 signaling in induction of VEGF-A expression, bone marrow–derived macrophages (BMMs) sorted for low Tie2 expression were transduced to overexpress a control (BMM/Tie2^Lo^) or Tie2 (BMM/Tie2^Hi^) (**Supplemental Figure 1A, 2A)**. Using these macrophages we found that Ang2 stimulation increased intracellular VEGF-A only in BMM/Tie2^Hi^ macrophages **(Supplemental Figure 1C, 1D)**. Furthermore, Tie2 knockdown in BMM/Tie2^Hi^ macrophages using Tie2 siRNA abrogated the ability of Ang2 to increase intracellular VEGF-A, confirming the requirement for Tie2 **(Supplemental Figure 2A-D; Figure 2F, G)**.

To relate these findings to our previous study showing that CSF-1 signaling drives VEGF-A secretion from macrophages without altering total VEGF-A expression ^15^, we pre-stimulated BAC1.2F5 macrophages with Ang2 in absence (Cont) or presence of Rebastinib (Reb) and then treated the cells with or without CSF-1 **(Figure 2H)**. As also demonstrated in result in Figure 2D, VEGF-A staining showed that Ang2 stimulation of macrophages increased intracellular VEGF-A, and this effect was blocked by rebastinib **(Figure 2I**, left panels; **Figure 2J**, left bars). Additional CSF-1 stimulation (+CSF-1) led to a significant decrease in intracellular VEGF-A compared with unstimulated cells (-CSF-1), with no change in VEGF-A levels in rebastinib-treated cells **(Figure 2I**, right panels; **Figure 2J**, right bars). We additionally found that Ang2 stimulation increased VEGF-A mRNA expression, and this induction was abrogated by rebastinib treatment **(Supplemental Figure 2C)**. Collecting conditioned media from these experiments, we found that pre-stimulation with Ang2 alone did not increase VEGF-A secretion **(Supplemental Figure 2D)**. In contrast, Ang2 pre-stimulation followed by CSF-1 treatment significantly enhanced VEGF-A secretion, an effect that was blocked by rebastinib **(Figure 2K**, gray vs. teal bars).

Taken together, these results indicate that Ang2/Tie2 signaling between endothelial cells and macrophages increases mRNA and intracellular protein expression of VEGF-A in macrophages, while CSF-1 signaling drives the subsequent release of this accumulated VEGF-A from macrophages.

### TMEM doorway macrophage VEGF-A expression is mediated by Tie2 signaling *in vivo*

We have previously shown that pharmacologic Tie2 inhibition with rebastinib *in vivo* reduces TMEM doorway associated vascular opening, circulating tumor cells (CTCs), and metastasis, and increases vascular integrity and overall survival in mice compared to control treatment ^30^. Given our *in vitro* findings that Tie2 activation increases VEGF-A expression in macrophages (**Figures 1-2**), we hypothesized that Tie2-dependent regulation of macrophage VEGF-A is the mechanism by which Tie2 signaling inhibition reduces TMEM doorway associated vascular opening and tumor cell dissemination. To investigate this, we analyzed primary tumors from mice treated with rebastinib or vehicle in two breast cancer metastasis models: the immunocompetent MMTV-PyMT syngeneic model **(Figure 3A-D)** and the HT17 patient-derived xenograft (PDX) model, in which patient tumors are orthotopically transplanted into *SCID* mice **(Figure 3E-H)**. Tumor sections were stained for VEGF-A and Iba1 to identify macrophages, and sequential sections were stained for TMEM doorways. As described in Figure 1A, sections were aligned at single-cell resolution, TMEM doorways were identified on the IHC slide, and VEGF-A expression was quantified specifically in TMEM doorway macrophages on the corresponding VEGF-A/Iba1 IF slide **(Figure 3B, 3F).** Tie2 inhibition significantly reduced intracellular VEGF-A levels in TMEM doorway macrophages **(Figure 3C, 3G)** and decreased extracellular VEGF-A in the pericellular region surrounding TMEM macrophages, across the entire TMEM doorway area, compared with vehicle-treated controls **(Figure 3D, 3H)**. These *in vivo* results are consistent with our *in vitro* data showing that blockade of Ang2/Tie2 signaling diminishes VEGF-A expression in macrophages (Figure 2), supporting a model in which Tie2 signaling maintains the elevated VEGF-A levels characteristic of TMEM doorway macrophages.

**Figure 3:**
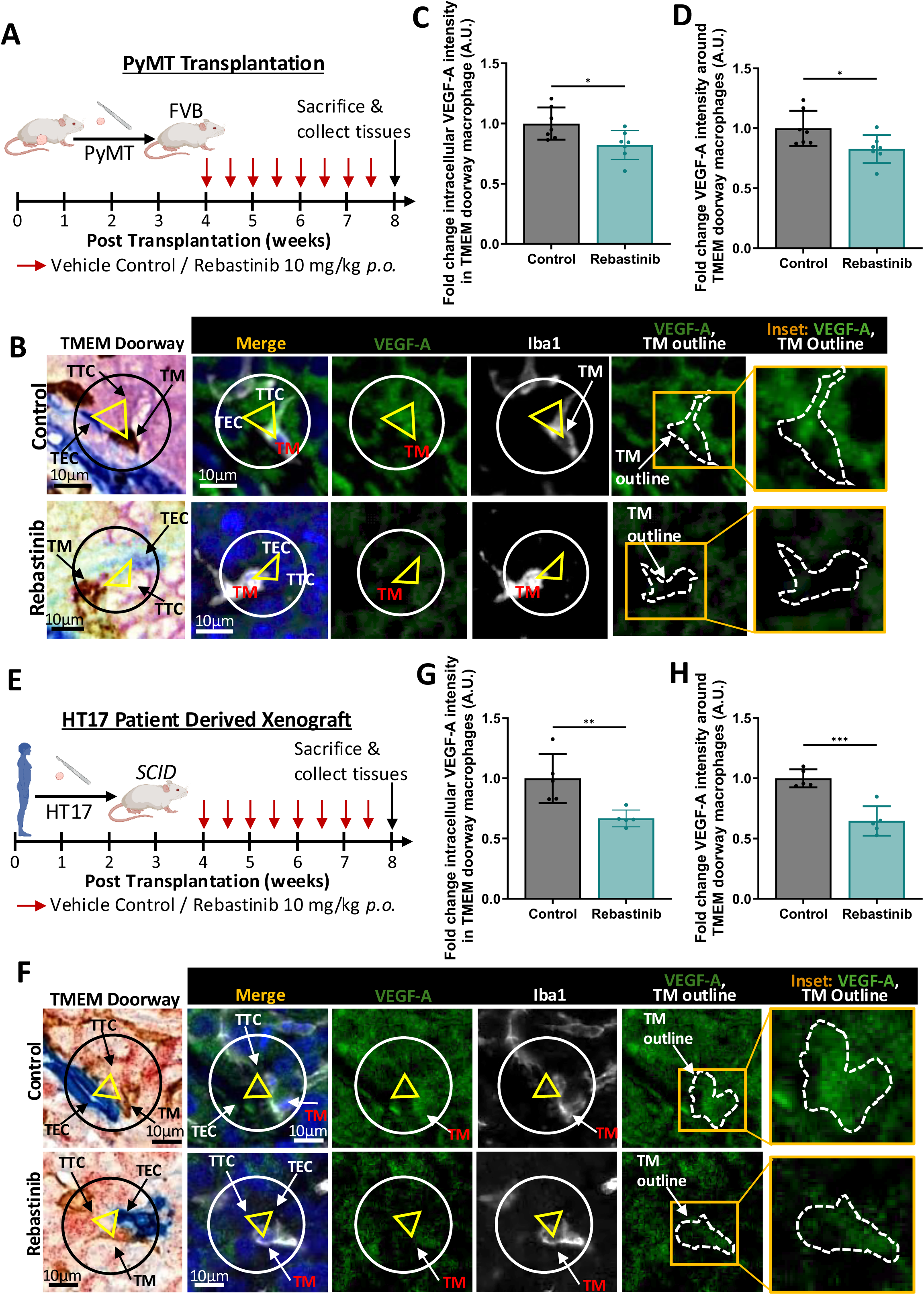
TMEM doorway macrophage VEGF-A expression is mediated by Tie2 signaling. **(A-D)** PyMT and **(E-H)** tumor chunks were orthotopically transplanted into (A-D) FVB and (E-H) *NOD-SCID* mice, respectively. Mice were treated twice a week for four weeks with 10 mg/kg Rebastinib or vehicle *p.o.* and primary tumors from these mice were used for staining. **(B, F)** Panels show immunostaining of VEGF-A intensity in TMEM doorway macrophages in tumor tissues from these mice treated with or without Rebastinib. Sequential tumor sections were stained by IHC (TMEM doorways-Mena, Iba1, endomucin) and immunofluorescence (VEGF-A (green), Iba1 (white), and DAPI (blue)). TMEM doorways were identified as described in Figure 1B. The circle in the IHC (black) and IF (white) panels show the same TMEM doorways obtained from the alignment of serial sections, and the three cells making up the TMEM doorway are indicated with the yellow triangle in each panel (TM, TEC, TTC). The TMEM doorway macrophage (TM) in the IF image, stained with Iba1, is enlarged in the rightmost panel. Scale bars= 20 µm. **(C-D, G-H)** The Iba1 (white) in the IF-stained slide within the TMEM doorway circle ROI was used to identify TMEM doorway macrophages and the immunofluorescence intensity of VEGF-A expression (green) within (intracellular, C, G) and outside (extracellular, D,H) the Iba1^+^ TMEM doorway macrophage was quantified. n= 5-7 mice per group, each dot represents the average value for a mouse. *p<0.05, **p<0.01, ***p<0.001, analyzed by Student’s *t*-test.

### Tie2 expressing macrophages are concentrated at and near TMEM doorways

While the *in vitro* and *in vivo* data indicate a clear role for Tie2-expressing macrophages in driving increased VEGF-A expression at TMEM doorways, multiple cell types in the tumor microenvironment express Tie2 and could be targeted by the Tie2 inhibitor, rebastinib. We therefore sought to identify which cells within the tumor microenvironment contribute to Tie2-dependent vascular opening at TMEM doorways and subsequent tumor cell dissemination. Using tissue staining and line-scan analysis of Tie2, together with cell lineage markers in mouse **(Figure 4A)** and human PDX **(Figure 4B)** breast cancer models, we detected Tie2 expression in endothelial cells, macrophages, and tumor cells. Although only ∼1% of Iba1^+^ macrophages expressed Tie2 when quantified across the entire tumor tissue **(Figure 4C)**, Tie2 expressing macrophages were located at an average of 2.49 μm from the closest TMEM doorway **(Figure 4D, 4E, Supplemental Figure 2E)** and 209.56 μm from the closest blood vessel without a TMEM doorway **(Figure 4D, 4F)**. Further, when focusing on perivascular regions and TMEM doorways specifically **(Figure 4G)**, 15% of perivascular macrophages expressed Tie2**, (Figure 4H)**. Of these Tie2-expressing macrophages, 90% were found within the 60 μm analysis regions surrounding TMEM doorways (defined by direct physical contact between a macrophage, tumor cell, and endothelial cell). **(Figure 4I)**. This pattern indicates that Tie2⁺ macrophages, while rare within the broader tumor microenvironment, are spatially enriched at TMEM doorways **(Figure 4I, 4J)**, providing a mechanism for selective VEGF-A accumulation at TMEM doorways where VEGF-A can be rapidly released to promote TMEM doorway opening and tumor cell intravasation.

**Figure 4:**
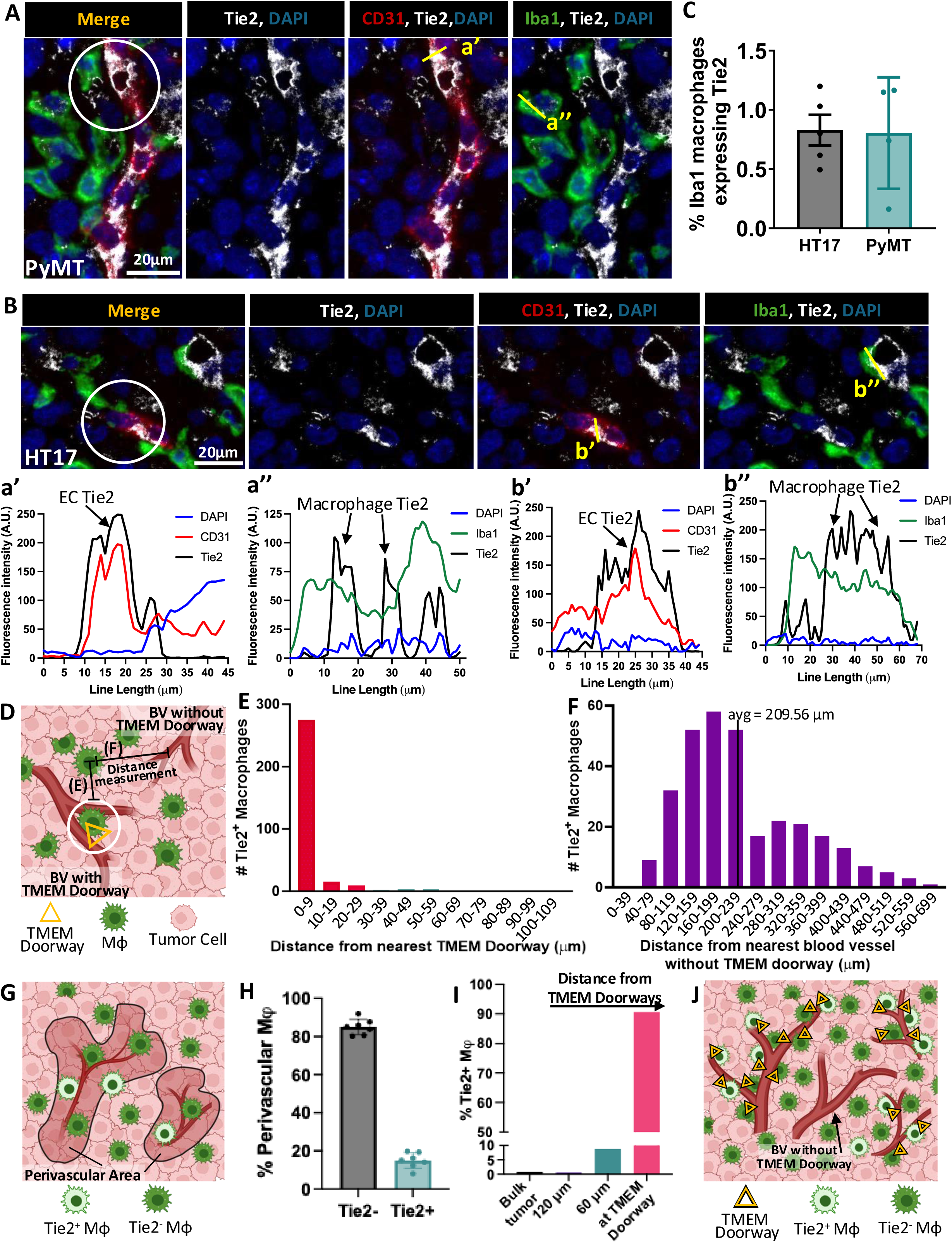
Tie2 expressing macrophages are concentrated near TMEM doorways. **A-B)** Tumor tissues from control mice in figure 3A (A, PyMT), and figure 3E (B, HT17), were stained using immunofluorescence (IF) with antibodies against CD31 (red), Iba1 (green), Tie2 (white). The TMEM doorways are indicated by the white circle. Scale bars= 20 µm. **a’-b”)** Quantification of the immunofluorescence intensity of Tie2, CD31 and Iba1 across the yellow line scans from IF stains in A and B. **C)** Quantification of the percentage of Tie2^+^/Iba1^+^ macrophages in PyMT and HT17 tumor tissue from A and B, using Visiopharm. n=4-5 mice per group, each dot represents the % of Iba1^+^ macrophages that express Tie2 for each mouse, +/-standard deviation. **D)** Quantification diagram of Tie2^+^ macrophage distance analysis from the nearest TMEM doorway (E) and from the nearest blood vessel without a TMEM doorway (F). White circle represents TMEM doorway and yellow triangle indicates cells of TMEM doorway. Diagram made using BioRender. **E)** Quantification of the distance of Tie2^+^/Iba1^+^ macrophages from TMEM doorways in PyMT and HT17 tumor sections stained in (A) and (B). Distances from 309 Tie2^+^/Iba1^+^ macrophages measured in 7 mice. Distances are binned every 10 μM, from 0 (at TMEM doorway) to 9 μM, 10 μm to 19 μM, 20 μm to 29 μm, etc. **F)** Quantification of the distance of Tie2^+^/Iba1^+^ macrophages from the nearest blood vessel without a TMEM doorways in HT17 tumor sections stained in (B). Distances from 309 Tie2^+^/Iba1^+^ macrophages measured in 7 mice. Distances are binned every 40 μM, from 0 (at TMEM doorway) to 39 μM, 40 μm to 79 μM, 80 μm to 119 μm, etc. **G)** Quantification diagram of the percentage Tie2^-^ macrophages (dark green cell) and Tie2^+^ macrophages (light green cell) in the perivascular space. The perivascular space was defined as within 20 μm of an endomucin stained blood vessel in HT17 tumor tissue from (B). Diagram made using BioRender. **H)** Quantification, as described in (G), of the percentage of perivascular macrophages that do (Tie2^+^) or do not (Tie2^-^) express Tie2 in PyMT and HT17 tumor tissue from (A) and (B), using Visiopharm. n=7 mice, each dot represents the average for one mouse, +/-standard deviation. **I)** Quantification of the percentage of Tie2^+^ macrophages in the tumor bulk (from C), and as the distance to TMEM doorways decreases. The right-most bar (pink) represents Tie2^+^ macrophages present in TMEM doorways. **J)** Cartoon summarizing the data found in panels E-I. Tie2^+^ macrophages are localized at perivascular regions, and more specifically at TMEM doorways and are not found near blood vessels without a TMEM doorway. Tie2^-^ macrophages represented with dark green cells and Tie2^+^ macrophages represented with light green cell. Yellow triangle represents a TMEM doorway. Diagram made using BioRender.

### Tie2 knockdown or inhibition *in vitro* blocks tumor cell transendothelial migration

To investigate the functional contribution of Tie2 signaling in each cell type to tumor cell intravasation, we used an established *in vitro* intravasation transendothelial migration (iTEM) assay, which models TMEM doorway conditions and allows quantification of tumor cell transendothelial migration **(Figure 5A)**. Prior work has shown that breast tumor cells seeded alone on matrigel with a confluent endothelial monolayer exhibit low basal transendothelial migration, which is significantly enhanced in the presence of macrophages ^42, 45^. To test the role of Tie2 signaling, tumor cells, endothelial cells, and macrophages were pretreated separately with the Tie2 inhibitor rebastinib or vehicle, washed, and then combined in the iTEM assay. As expected, tumor cells displayed a basal level of transendothelial migration that significantly increased when macrophages were present **(Figure 5B–D**, first two bars). Rebastinib pretreatment of endothelial cells (ECs) or tumor cells did not significantly alter tumor cell transendothelial migration **(Figure 5B, 5C**, last bar), whereas rebastinib pretreatment of macrophages (Mφ) significantly reduced tumor cell transendothelial migration **(Figure 5D**, last bar). These findings suggest that Tie2 signaling in macrophages, but not in tumor or endothelial cells, is necessary for macrophage-driven enhancement of transendothelial migration.

**Figure 5:**
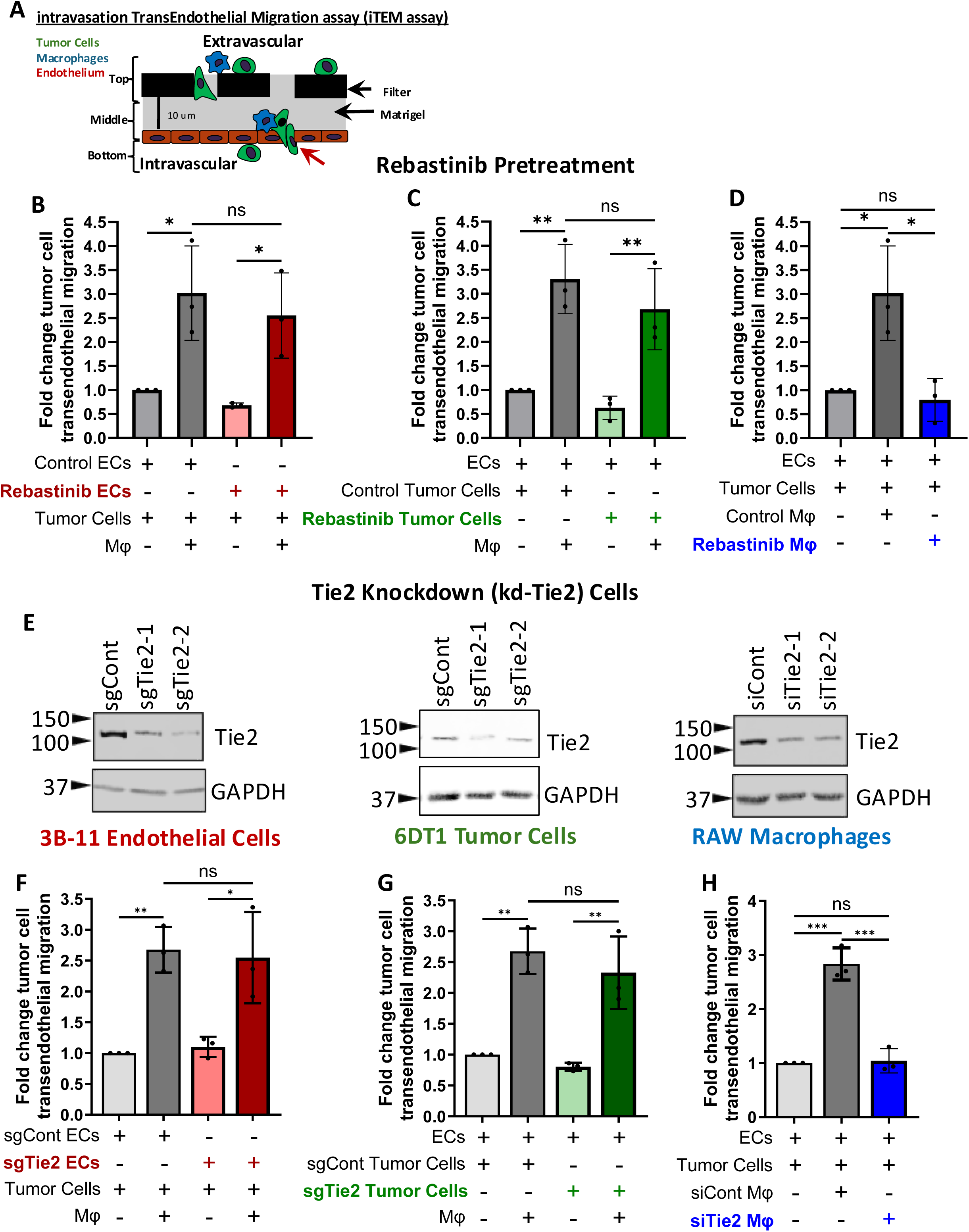
Tie2 knockdown or inhibition in macrophages *in vitro* blocks tumor cell transendothelial migration. **A)** Diagram of subluminal to luminal intravasation trans-endothelial migration (iTEM) assay, which is a model to measure ability of tumor cells to intravasate. Tumor cells (6DT1, green) cells are labeled with CellTracker^TM^ green and plated on a confluent endothelium (3B-11 cells, brown) either alone or with RAW macrophages labeled with CellTracker FarRed. After 18 hours incubation, the cells are fixed and the endothelium is stained with ZO-1 antibodies to ensure endothelial confluence and tight junctions at locations of TC crossing. The number of green 6DT1 tumor cells which cross the endothelium are imaged using confocal microscopy and quantified using ImageJ software. **B-D)** 3B-11 endothelial cells (B), 6DT1 tumor cells (C), and RAW macrophages (D), were pretreated with 1 nM Rebastinib or vehicle control overnight and the next day, cells were washed and combined in the iTEM assay in the combinations shown in the figure. n=3 experiments, each point represents the average fold change tumor cell trans-endothelial migration compared to control ECs and tumor cells alone (first bars in each graph). *p<0.05, **p<0.01, analyzed by one-way ANOVA. **E)** Western blot of lysates from Control (sgCont, siCont) and Tie2 knockdown (sgTie2, siTie2) 3B-11 endothelial cells (ECs), 6DT1 tumor cells, and RAW macrophages (M⏀) using antibodies against Tie2 and GAPDH. Representative blots shown from n=3 experiments. **F-H)** Tie2 knockdown and control 3B-11 endothelial cells (F), 6DT1 tumor cells (G), and RAW macrophages (H), from (E) were combined in the iTEM assay in the combinations shown in the figure. n=3 experiments, each point represents the average fold change tumor cell trans-endothelial migration compared to control ECs and tumor cells alone (first bars in each graph). *p<0.05, **p<0.01, ***p<0.001, analyzed by one-way ANOVA.

As a secondary approach, Tie2 expression was reduced using CRISPR-Cas9-mediated knockdown in endothelial and tumor cells (sgTie2 vs sgCont) and siRNA-mediated knockdown in macrophages (siTie2 vs siCont) **(Figure 5E)**. While control and wild-type macrophages again promoted tumor cell transendothelial migration **(Figure 5F-H**, first two bars), Tie2 knockdown in macrophages significantly decreased it **(Figure 5H**, last bar). Importantly, Tie2 knockdown in endothelial or tumor cells did not significantly affect tumor cell transendothelial migration relative to controls **(Figure 5F, 5G,** last two bars), reinforcing that, although tumor cells, endothelial cells, and macrophages all express Tie2, macrophage Tie2 is specifically required to support tumor cell transmigration across the endothelium.

### Tamoxifen-induced Cre activation selectively depletes Tie2 in tumor-associated macrophages

To test the role of Tie2-expressing macrophages *in vivo,* we generated an inducible macrophage-specific Tie2 knockout mouse by floxing the first exon of *Tek* (Tie2^fl/fl^) **(Supplemental Figure 3)** and crossing these mice with animals expressing a tamoxifen-inducible Cre recombinase under the control of the macrophage specific Csf1r promoter (tg*Csf1r-Mer-iCre-Mer*) ^35^. Isolated bone marrow-derived macrophages from Cre^+^ mice treated with tamoxifen (+Tam) showed significant reduction of Tie2 protein expression compared to Cre^-^ and vehicle control (-Tam) treated mice **(Figure 6A)**. Within the tumor microenvironment, tamoxifen-induced Cre activity most efficiently targets tumor associated macrophages, which are long-lived and exhibit strong CSF-1R expression, followed by monocytes and monocytic MDSCs, which also express CSF-1R. With prolonged tamoxifen exposure, heterogeneous recombination can occur in dendritic cells and neutrophils as well. Because this broader myeloid targeting could confound interpretation, a tamoxifen time-course experiment was performed to determine the optimal tamoxifen treatment time to selectively target Tie2 in macrophages while sparing other cells within the tumor microenvironment **(Supplemental Figure 4A)**. Two doses of tamoxifen at 72 and 96 hours before sacrifice significantly reduced Tie2 expression in intratumoral macrophages **(Supplemental Figure 4B, C)**, without significantly altering Tie2 expression in endothelial cells, monocytes, MDSCs, neutrophils, dendritic cells, or tumor cells **(Supplemental Figure 4D-I)**.

**Figure 6:**
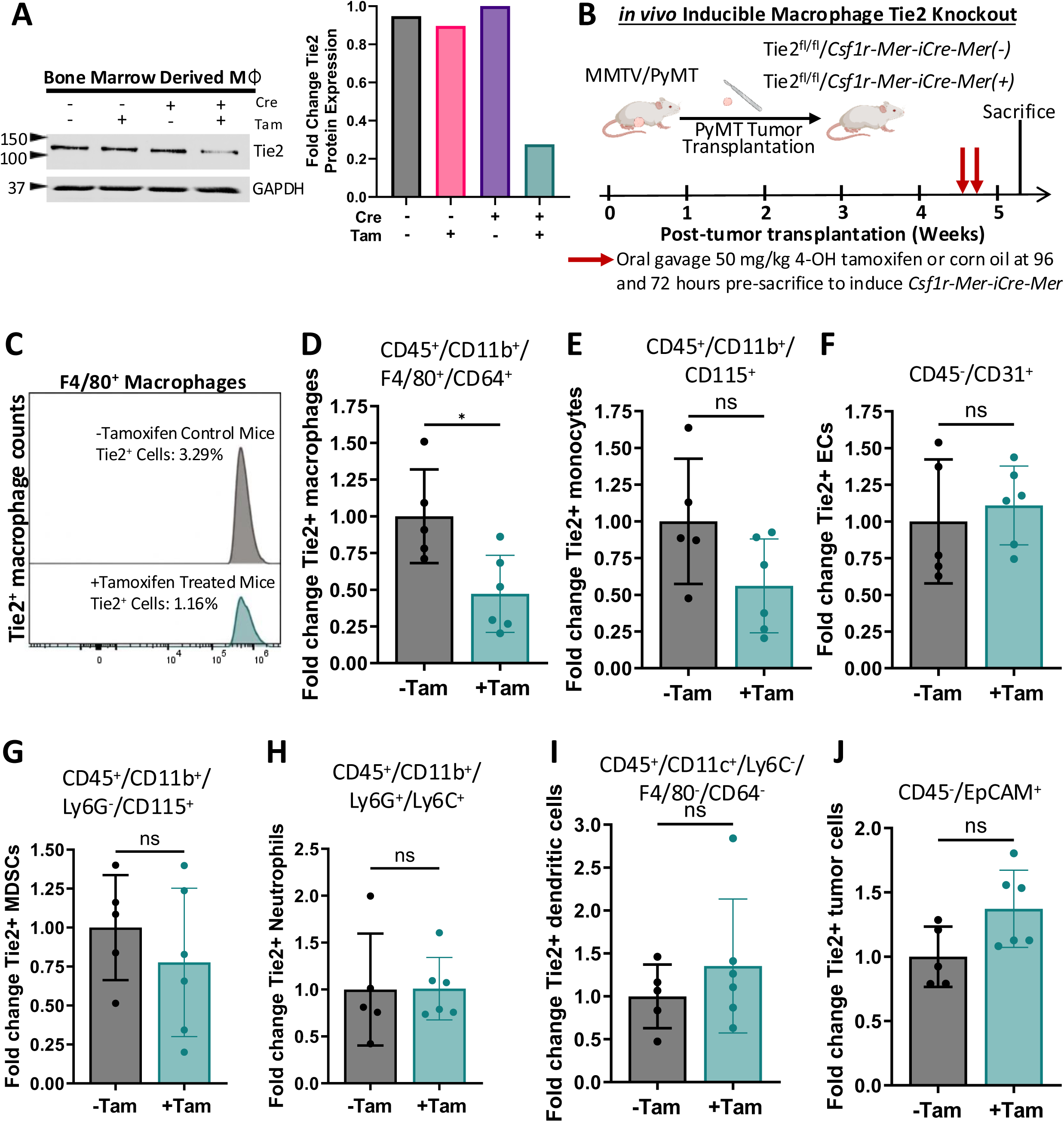
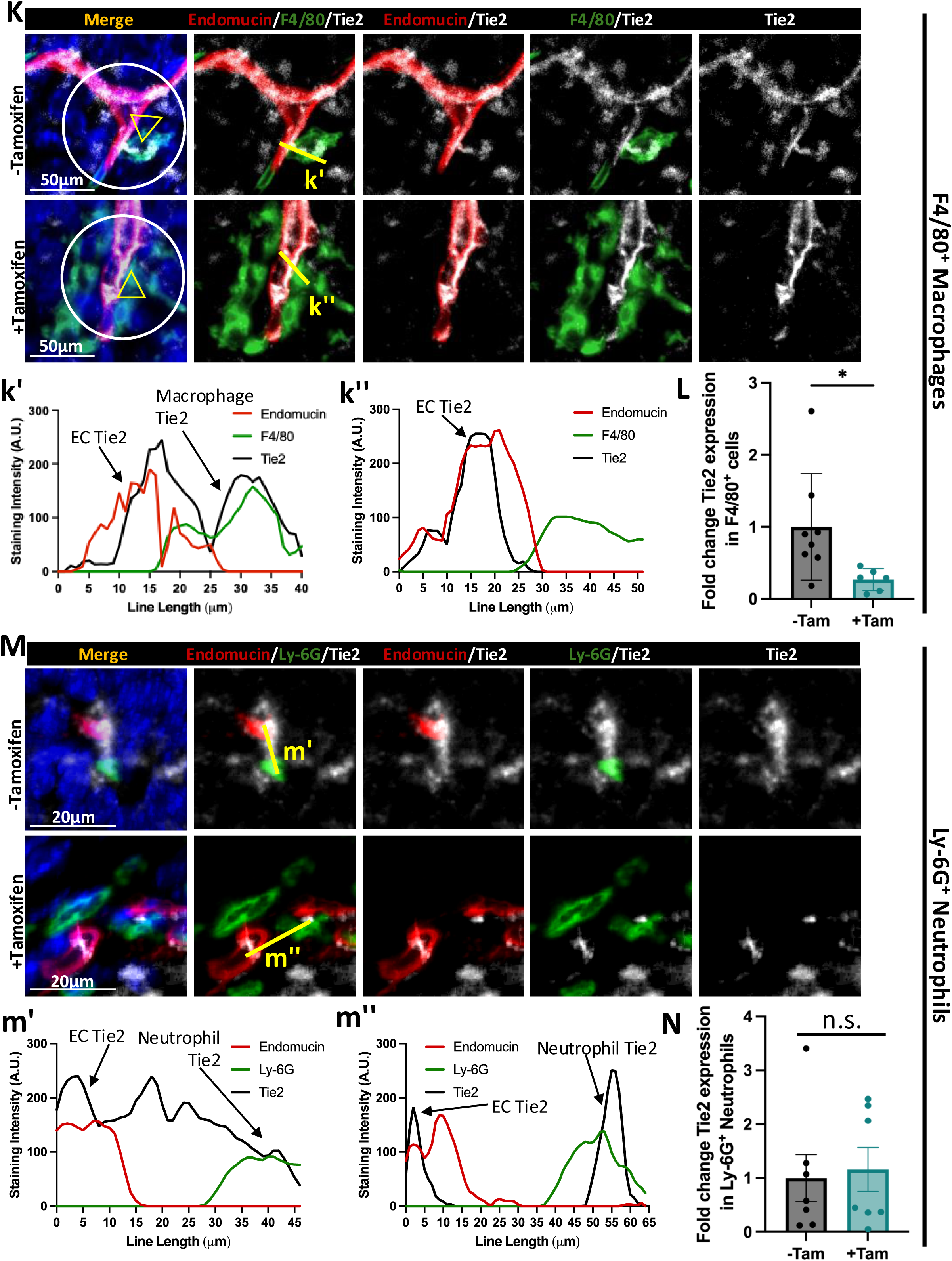
Tie2 is knocked out macrophages in the tumor microenvironment. **A)** Bone marrow macrophages isolated from Tie2^fl/fl^/*Csf1r-Mer-iCre-Mer*^+^ (Cre+) and Tie2^fl/fl^/*Csf1r-Mer-iCre-Mer* ^−^ (Cre-) mice were treated with 50 mg/kg 4-OH tamoxifen (+Tam) or corn oil (-Tam) by oral gavage, 96 and 72 hours before sacrifice. Lysates were used in a western blot using antibodies against Tie2 and GAPDH. Graph shows quantification of fold change in Tie2 protein expression in bone marrow macrophages from western blot, normalized to GAPDH loading control and set relative to +Cre, -Tam treated mice. **B)** Experimental model of inducible, conditional knockout of Tie2 in macrophages. Tie2^fl/fl^/*Csf1r-Mer-iCre-Mer*^+^ (Cre+) and Tie2^fl/fl^/*Csf1r-Mer-iCre-Mer* ^−^ mice were treated with 50 mg/kg of 4-OH tamoxifen (+Tam) or corn oil (-Tam) by oral gavage, 96 and 72 hours before sacrifice. Fifteen minutes before sacrifice, mice were injected *i.v.* with 20mg/mL 155kDa TMR-dextran. Blood was removed by cardiac puncture for CTC quantification. Primary tumors and lungs were removed and digested for flow cytometry and MACS experiments or fixed for staining experiments. **C)** Tie2^+^ macrophage counts from parental macrophage population (CD45^+^CD11b^+^F4/80^+^CD64^+^) in mice treated with corn oil (-Tam, gray population) or 4-OH tamoxifen (+Tam, teal population). **D-J)** Graphs depict fold change in Tie2^+^ D) macrophages, E) monocytes, F) endothelial cells, G) myeloid derived suppressor cells (MDSCs), H) neutrophils, I) dendritic cells, and J) tumor cells compared to control mice (+Cre, -Tam) from mice treated with corn oil (-Tam) or tamoxifen (+Tam) as described in (C). **K, M)** Primary tumor sections from mice from (C) were stained with for antibodies against Tie2, F4/80, PyMT, Ly-6G, and endomucin. TMEM doorways are localized within the white circle and indicated with the yellow triangle in the merge panel. k’ and m’ line scans are along the TMEM doorway endothelial cell and macrophage in control (-Tamoxifen) treated mice. k” and m” line scans are along TMEM doorway endothelial cell and macrophage from Tamoxifen treated mice. **(m’-d”)** Quantification of the immunofluorescence intensity of (k’-k”) Tie2, endomucin and F4/80 and (m’-m”) Tie2, endomucin, and Ly-6G across the yellow line scan from “Merge” in (K and M). **L)** Quantification of % of Tie2+/F4/80+ Tie2 macrophages in tumor sections from mice treated in (C). **N)** Quantification of % of Tie2+/Ly-6G+ Tie2 neutrophils in tumor sections from mice treated in (C).

To test the effects of acute, macrophage-specific Tie2 blockade in established tumors on TMEM doorway associated vascular opening and tumor cell dissemination, tumors from MMTV-PyMT mice were orthotopically transplanted into Tie2^fl/fl^/Csf1r-Mer-iCre-Mer^-^ and Tie2^fl/fl^/Csf1r-Mer-iCre-Mer^+^ recipient mice. Once the tumors reached at least 0.6 cm^3^ in volume, mice were treated with corn oil or tamoxifen to induce macrophage specific knockdown of Tie2. **(Figure 6B)**. Flow cytometric analysis of tumors showed a ∼53% reduction in Tie2^+^ macrophages (CD45^+^/CD11b^+^/F4/80^+^/CD64^+^) from tamoxifen-treated mice compared with controls (**Figure 6C, 6D**), with no significant knockdown of Tie2 in other myeloid populations potentially targeted by Cre activation, including MDSCs, neutrophils, and dendritic cells **(Figure 6E-I, Supplemental Figure 5)**. As expected, this treatment did not alter Tie2 expression in tumor cells or endothelial cells, the other cellular components of TMEM doorways **(Figure 6F, 6J, Supplemental Figure 5)**. Consistent with these findings, tissue staining demonstrated an approximately 73% decrease in Tie2^+^ F4/80^+^ macrophages in tamoxifen-treated tumors **(Figure 6K, 6L).** Line-scan analysis showed Tie2 expression in TMEM doorway endothelial cells was maintained in both treatment groups, whereas Tie2 expression in TMEM doorway macrophages was selectively reduced in tamoxifen-treated animals compared to controls **(Figure 6k’-k”)**. As neutrophils are known contributors to breast cancer metastasis ^46^ and Tie2^+^ neutrophils could also be targeted in this model, we specifically examined Tie2 expression in neutrophils. In agreement with the flow cytometry analysis (**Figure 6H**), tissue staining and line scan analysis revealed no change in Tie2 expression in Ly-6G^+^ neutrophils between treatment groups **(Figure 6M, 6N, 6m’-m”, 6n’-n”)**. Interestingly Ly-6G^+^ neutrophils were relatively rare compared to F4/80^+^ macrophages in this tumor model **(Supplemental Figure 6A-D)**. Altogether, we are confident that this mouse model selectively and effectively targets Tie2 expression in macrophages within the tumor microenvironment.

### Inducible knockdown of Tie2 in macrophages *in vivo* decreases metastatic dissemination and TMEM doorway activity

To examine the effects of macrophage-specific Tie2 knockdown on TMEM doorway-associated vascular opening, one cohort of mice was injected with high-molecular weight TMR-dextran 15 minutes before sacrifice and tumor sections were stained for endomucin and TMR-dextran **(Figure 7A)**. Specific targeting of Tie2^+^ macrophages with tamoxifen (+Tam) significantly decreased TMEM doorway-associated vascular opening, as dextran extravasation was inhibited and was retained within the vasculature **(Figure 7B)**. Co-localization of the endothelial adherens junction protein CD31 and tight junction protein ZO-1 increased in tamoxifen treated mice compared to controls, indicating reduced endothelial permeability following macrophage-specific Tie2 knockdown **(Figure 7C, D)**.

**Figure 7:**
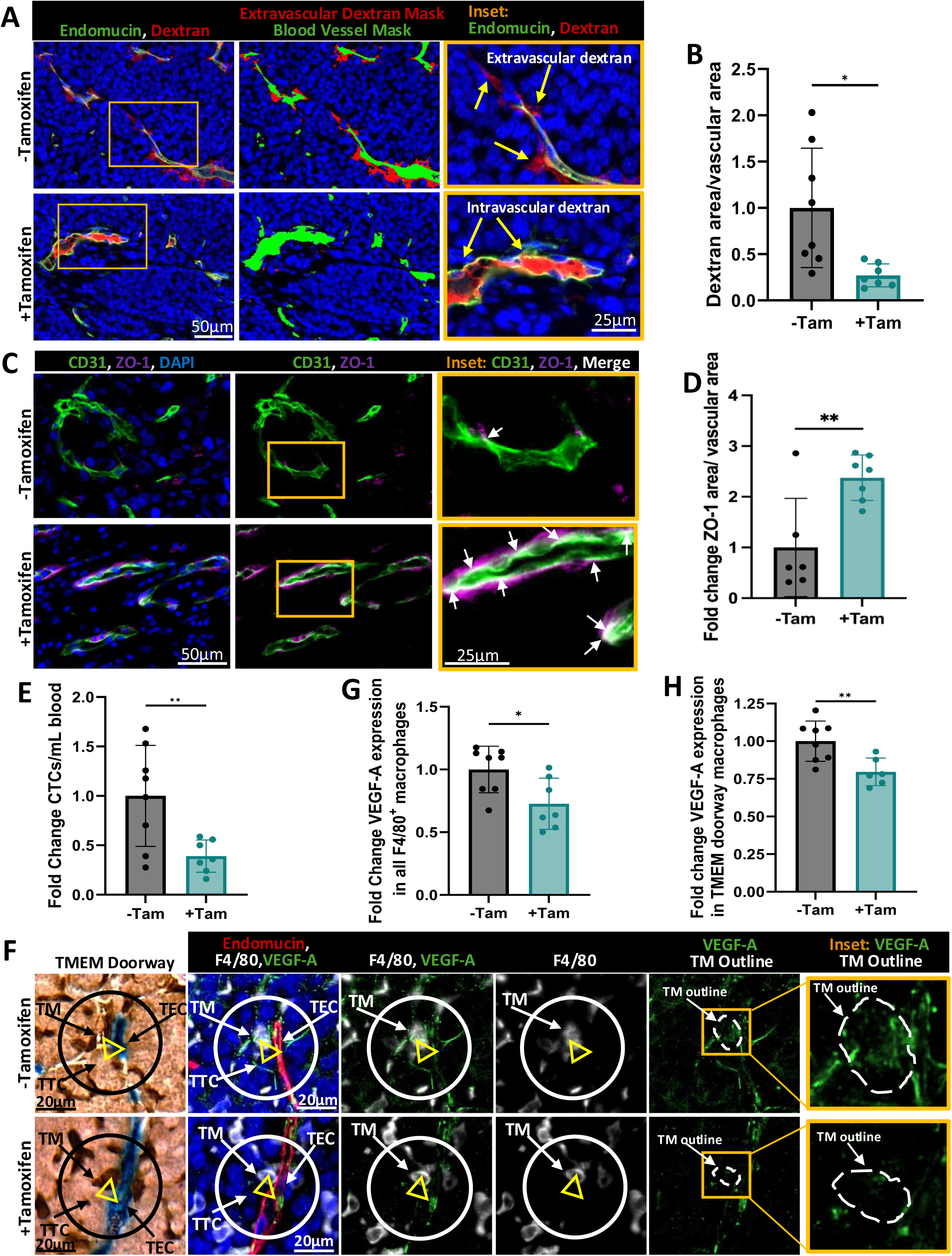
Inducible knockout of Tie2 in macrophages *in vivo* decreases metastatic dissemination and TMEM doorway activity. **A)** Primary tumors from figure 6C were stained with endomucin (green), TMR-dextran (red), and DAPI (blue). Middle panel shows the blood vessel mask from the endomucin stain (green) and the extravascular dextran mask (red). Right panel shows increased magnification of yellow box from the left panel to demonstrate the localization of dextran in relation to the vessel lumen. Left and middle panels scale bar=50 µm; left panel= 25 µm. **B)** Quantification of extravascular 155 kDa TMR-dextran. *p<0.05, analyzed by Student’s *t*-test, n=7-8 mice per group, each dot represents the average value for a mouse. **C)** Mice from (A) stained with CD31 (green), ZO-1 (magenta), and DAPI (blue). Right panel shows increased magnification of yellow box from middle panel to demonstrate overlap between CD31 and ZO-1 stains (merge/overlap of the two stains indicated with white signal and indicated with white arrows). Left and middle panels scale bar=50 µm; left panel= 25 µm. **D)** Quantification of vascular ZO-1 stains in (C), **p<0.01 analyzed by Student’s *t*-test, n=7-8 mice per group, each dot represents the average value for a mouse. **E)** Quantification of circulating tumor cells from mice in Figure 6C. **p<0.01, analyzed by Student’s *t*-test, n=7-8 mice per group, each dot represents the normalized value for a mouse. **F)** Immunostaining of VEGF-A intensity in TMEM doorway macrophages, obtained from mice in (A). Sequential tumor sections were stained by IHC (TMEM doorways-Mena, Iba1, endomucin) and immunofluorescence (IF, VEGF-A (green), F4/80 (white), endomucin (red), and DAPI (blue)). TMEM doorways were identified as described in Figure 1A. The circle in the IHC (black) and IF (white) panels show the same TMEM doorways obtained from the alignment of serial sections, and the three cells making up the TMEM doorway are indicated with the yellow triangle in each panel (TM, TEC, TTC). The TMEM doorway macrophage (TM) in the IF image is stained with F4/80 (white) and is outlined in white in the VEGF-A IF panel. Right panel shows increased magnification of yellow box from the VEGF-A IF panel, showing VEGF-A expression within the TMEM doorway macrophage. Scale bars= 20 µm. **G)** Quantification of VEGF-A expression in F4/80^+^ macrophages from mice in treatment regimen illustrated in Figure 6C. n= 7-8 mice per group, each dot represents the average value for a mouse. *p<0.05, analyzed by Student’s *t*-test. **H)** The F4/80 stain (white) in the IF-stained slide within the TMEM doorway circle ROI was used to identify TMEM doorway macrophages and the immunofluorescence intensity of VEGF-A expression (green) within the F4/80 TMEM doorway macrophage was quantified. n= 7-8 mice per group, each dot represents the average value for a mouse. **p<0.01, analyzed by Student’s *t*-test.

At the time of sacrifice, we collected blood from the mice to quantify circulating tumor cells (CTCs) and found that Tie2 knockdown in macrophages (+Tam) significantly reduced the number of CTCs compared to control mice (-Tam) **(Figure 7E)**. To analyze VEGF-A expression in macrophages we stained sequential tumor sections for TMEM doorways (IHC stain; **Figure 7F)** and for endomucin, F4/80, and VEGF-A (IF stain, **Figure 7F)**. We found an overall decrease in VEGF-A expression in F4/80^+^ macrophages in Tie2-macrophage knockdown mice, both when measuring across the tumor bulk **(Figure 7G)** and specifically at TMEM doorway macrophages **(Figure 7H)**, consistent with Ang2/Tie2 signaling sustaining increased VEGF-A expression in TMEM doorway macrophages. We further found that treatment of PyMT tumor bearing mice with rebastinib plus paclitaxel significantly decreased the paclitaxel-induced induction of CTCs compared to paclitaxel alone treated mice **(Supplemental Figure 7A-B)**, demonstrating that inhibition of Tie2 signaling inhibits tumor cell dissemination. These *in vivo* findings align with the *in vitro* data showing that macrophage specific Tie2 inhibition decreases tumor cell transendothelial migration (Figure 5) and VEGF-A expression in macrophages (Figure 2), supporting a model in which Ang2/Tie2 signaling drives the accumulation of VEGF-A in perivascular macrophages at TMEM doorways. Upon subsequent CSF-1 stimulation, this locally stored VEGF-A is released to trigger precise opening of TMEM doorways, thereby facilitating tumor cell intravasation and dissemination **(Figure 8)**.

**Figure 8:**
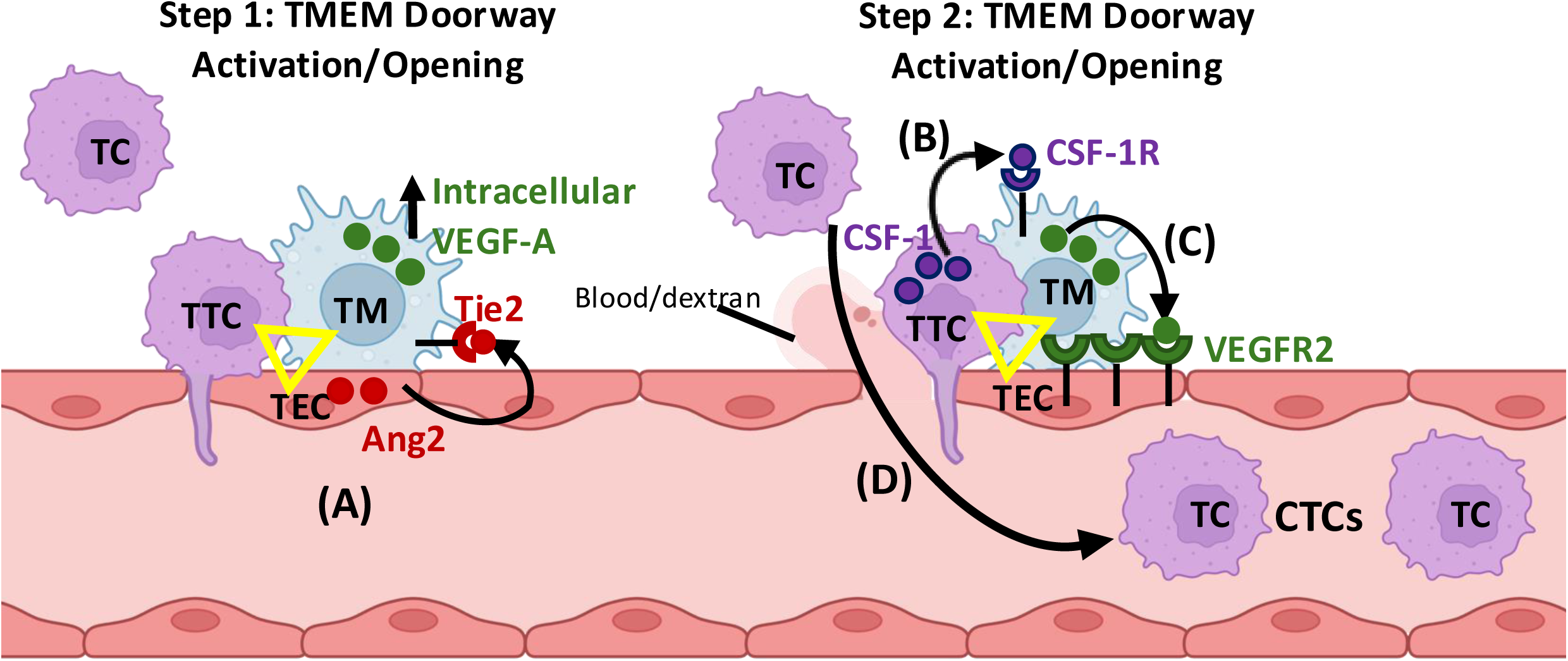
Mechanism of TMEM doorway opening and tumor cell dissemination. The TMEM doorways are denoted with yellow triangles, with each point of the triangle at a TMEM doorway tumor cell (TTC), endothelial cell (TEC), or macrophage (TM). TMEM doorway cells are in direct and stable physical contact throughout the TMEM doorway activation or opening process ^1, 3, 4^. **A)** In step 1 of TMEM doorway activation, TECs secrete Ang2 which binds to the Tie2 receptor, expressed on TMs. Activation of Ang2/Tie2 signaling causes TMs to increase expression of VEGF-A mRNA and protein; however, VEGF-A protein is not secreted. Critically, Tie2 expression is required for upregulation of VEGF-A within perivascular macrophages. **B)** In step 2 of TMEM doorway activation/opening, CSF-1 secreted by TTCs binds to the CSF-1R receptor on TMs, triggering the local secretion of VEGF-A within the TMEM doorway. **C)** VEGF-A binds to VEGFR2, expressed on TECs, leading to TMEM doorway-associated vascular opening. This is an acute event, with the vascular opening only occurring at the TMEM doorway and lasting ∼20 minutes before the vasculature at the TMEM doorway is resealed. **D)** While the TMEM doorway is opened and active, vascular contents extravasate (e.g. blood/dextran) and streaming tumor cells intravasate through the opening, forming CTCs. The left TMEM doorway is inactive, as there is no vascular opening, and the right TMEM doorway is active, as the vasculature is open, allowing movement of cells and contents between the vasculature and tissue. Inhibition or knockdown of Tie2 specifically in macrophages prevents Ang2-induced VEGF-A expression. Without the accumulation of VEGF-A within the TMEM doorway macrophage, the TMEM doorway cannot be signaled to open, and tumor cells cannot intravasate into the vasculature.

## Discussion

Here, we investigated how endothelial cell-derived Ang2 and macrophage Tie2 signaling cooperate to control VEGF-A expression in TMEM doorway macrophages, TMEM doorway opening, and metastatic dissemination in breast cancer. We have previously shown using intravital imaging that vascular opening associated with metastatic dissemination in tumors is an acute, localized, and transient event that occurs exclusively at TMEM doorways, in contrast to intravenous injection of VEGF-A, which induces generalized and continuous vessel leakage ^1^. We further demonstrated that macrophage specific VEGF-A knockout inhibited both TMEM doorway opening and tumor cell intravasation, underscoring the need to understand the upstream mechanisms that drive VEGF-A accumulation in TMEM doorway macrophages, especially given the clinical failure of anti-VEGF-A and anti-angiogenic therapies to prevent metastasis^1, 47^. These therapies are thought to be limited by tumor evasion mechanisms such as upregulation of alternative pro-angiogenic factors, including angiopoietins, recruitment of bone marrow-derived proangiogenic monocytes/macrophages, and enhanced tumor cell invasion and metastasis^47^. Critically, inhibition of Tie2 signaling in macrophages at TMEM doorways could address all of these modes of anti-angiogenic therapy resistance.

Using multiplex staining and dextran-based reporters of vascular opening in mouse models, we found that Ang2 accumulates extravascularly at active TMEM doorways, where vascular opening and tumor cell intravasation occur. *In vitro* assays demonstrated that mechanistically, endothelial cells, via Ang2/Tie2 paracrine signaling, selectively increase VEGF-A mRNA and intracellular protein in Tie2⁺ macrophages, while CSF-1 signaling triggers secretion of this stored VEGF-A^15^. Complementary pharmacologic and genetic Tie2 inhibition in *in vitro* models of tumor cell intravasation revealed that Ang2/Tie2 signaling in macrophages, but not in tumor cells or endothelial cells, is necessary for macrophage-driven tumor cell trans-endothelial migration.

Extending these findings *in vivo*, we utilized pharmacologic Tie2 inhibition and an inducible, macrophage-specific conditional Tie2 knockout model to dissect the role of macrophage Tie2 in established tumors. Tie2 blockade or macrophage-specific Tie2 deletion reduced VEGF-A expression in TMEM doorway macrophages, enhanced endothelial junction integrity at TMEM doorways, inhibited TMEM doorway-associated vascular openings, and decreased the levels of CTCs and hence metastatic dissemination. Spatial mapping of Tie2^+^/F4/80^+^ macrophages revealed that Tie2⁺ macrophages, although rare in the overall tumor, as previously reported ^30, 48^, are preferentially concentrated at and near TMEM doorways. This creates perivascular VEGF-A “hotspots” that, upon CSF-1 stimulation, can drive localized vessel opening and subsequent tumor cell intravasation and dissemination to secondary sites ^1, 2, 49^.

Altogether, these findings define an Ang2/Tie2—CSF-1 signaling axis between endothelial cells, TMEM doorway macrophages, and tumor cells that focuses VEGF-A-dependent vascular permeability to discrete vascular openings for intravasation. This two-step model resolves several key paradoxes: (1) why VEGF-A secretion-mediated vascular opening occurs selectively at TMEM doorways ^1^ despite ubiquitous TAM presence in the tumor, (2) why CSF-1/CSF-1R inhibition abrogates metastasis without eliminating TMEM doorway macrophage VEGF-A expression ^15^, and (3) how the rare population of Tie2^+^ macrophages, beyond induction of tumor neoangiogenesis ^24, 27^, exerts an outsized impact on metastasis via their spatial concentration at TMEM doorways. These findings offer a rationale for selectively targeting Tie2-expressing macrophages to inhibit breast cancer dissemination in patients.

TMEM doorways represent the sole sites of tumor cell intravasation in breast cancer and the TMEM doorway score within primary tumors serves as a clinically validated prognostic marker of distant metastatic recurrence in treatment naïve patients with ER^+^/HER2^-^ breast cancer ^3–5^. This prognostic value extends to patients with residual ER^+^/HER2^-^ breast cancer after neoadjuvant chemotherapy ^50^, where TMEM doorway score is further increased by neoadjuvant chemotherapy itself ^37, 51^. The clinical relevance of TMEM doorways extends further to racial disparities in breast cancer outcomes: Black patients not only have higher TMEM doorway score in their ER^+^/HER2^-^breast cancer compared to White patients ^6, 50^ but are also 4.6-fold more likely to have distant recurrence if they have high TMEM doorway score compared to white patients ^6^, partially explaining the racial disparities seen in ER^+^/HER2^-^ disease ^52–54^.

Importantly, the therapeutic potential of targeting this pathway has begun to be tested clinically: a recently completed phase 1b clinical trial demonstrated the safety of the Tie2 inhibitor rebastinib combined with chemotherapeutic agents, paclitaxel or eribulin in patients with HER2^-^metastatic breast cancer ^55^. Patients treated with rebastinib and chemotherapy saw a significant decrease in CTCs and circulating Ang2 levels following two cycles of therapy, and the therapy was well tolerated ^55^. The mechanistic framework established here thus provides a molecular basis for the prognostic power of TMEM doorway score across these patient populations and underscores the rationale for targeting Tie2-expressing macrophages as an antimetastatic strategy.

Tie2-expressing macrophages associate with endothelial cells to drive tumor angiogenesis during tumor progression ^24^ and promote vascular reconstruction following chemotherapy treatment ^27^. We previously showed that the Tie2 inhibitor rebastinib reduces metastasis in orthotopic breast cancer mouse models ^30, 37^. However, a recent study (Jakeb et al.) using a constitutive myeloid Tie2 knockout model (*LysM-Cre*) reported that Tie2^+^ macrophages are not required for tumor growth, revascularization, and metastasis ^48^. Notably, the authors came to this conclusion by using ectopic model: examining the lung metastasis of subcutaneously injected Lewis Lung Carcinoma cells. Furthermore, by only comparing chemotherapy alone versus chemotherapy plus Tie2 inhibition, without a vehicle control group, the study could not assess the effects of Tie2 inhibition on metastasis independently of chemotherapy ^48^.

In our previous studies using orthotopic breast cancer metastasis models, we similarly observed no difference in lung metastatic foci between chemotherapy alone versus chemotherapy plus rebastinib ^30^. However, rebastinib alone or combined with chemotherapy significantly reduced lung metastatic foci when compared to vehicle controls ^30^. Additionally, we found that rebastinib combined with the chemotherapeutic eribulin significantly improved overall survival in mice with metastatic breast cancer compared to both eribulin-alone and control-treated mice ^30^. In this current study, using the first-line chemotherapy, paclitaxel, we found that rebastinib combined with paclitaxel significantly decreased CTCs in primary tumor bearing mice compared to mice treated with paclitaxel alone (Supplemental Figure 7), underscoring that inhibition of Tie2 at TMEM doorways decreases tumor cell intravasation and formation of CTCs.

Using immunofluorescence staining and flow cytometry in several models (Figure 4C, 6D), we found that Tie2+ macrophages represent a rare population within the tumor, consistent with Jakab et al., who made the same observation using scRNAseq data in human and mouse tumors^48^. However, despite this rarity, our current results show that Tie2^+^ macrophages are spatially enriched at TMEM doorways, the defined sites of tumor cell intravasation.

Co-staining Tie2 with multiple endothelial cell and macrophage markers confirmed that we were identifying Tie2^+^ macrophages and excluding endothelial cells. We found that a subset of perivascular macrophages expressed Tie2, and that the majority of Tie2^-^expressing macrophages were localized at TMEM doorways (Figure 4H, 4E). This selective perivascular enrichment of Tie2^+^ macrophages at TMEM doorways provides a mechanism for localized VEGF-A accumulation and rapid release at defined vascular hotspots, enabling acute and confined TMEM doorway opening and tumor cell intravasation observed by intravital imaging ^1^.

Our study also defines a new mouse model and protocol for acute targeting of Tie2-expressing macrophages within the tumor microenvironment (**Figure 6, Supplemental Figure 4**), avoiding the promiscuity which plagues myeloid-driving Cre mouse models ^56^. Here we utilized the tamoxifen-inducible *Csf1r-Mer-iCre-Mer* Cre driving mouse, which employs an improved cre (iCre) and preferentially recombines long-lived, CSF-1R-high tumor-associated macrophages over monocytes/neutrophils ^35^. Tamoxifen and 4-OH tamoxifen dosage and time-course optimization achieved ∼50-70% Tie2 reduction in tumor macrophages while sparing monocytes, MDSCs, neutrophils, dendritic cells, and endothelial cells, as validated by flow cytometry, immunofluorescence staining of cell lineage markers, and line scan analysis. While this model is ideal for short term studies of Tie2 inhibition, longer term use of tamoxifen will cause broad targeting of Tie2 within the myeloid lineage and confound interpretation of macrophage/monocyte specific studies. This precision targeting reveals the specialized role of Tie2^+^ macrophages in TMEM doorway-mediated tumor cell intravasation and dissemination.

Here, we provide the first complete, spatially resolved signaling mechanism for TMEM doorway function, identifying the Ang2/Tie2-CSF-1-VEGF-A paracrine signaling network as the core regulatory mechanism governing metastatic intravasation. Three novel advances emerge: (1) spatial logic explaining restriction of VEGF-A release to TMEM doorways sites through Tie2^+^ macrophage enrichment; (2) division of labor between Ang2/Tie2 (VEGF-A expression) and CSF-1 (VEGF-A release); and (3) cell-type specificity demonstrating that macrophage, but not endothelial or tumor cell, Tie2 mediates TMEM doorway function. By establishing spatial enrichment, cell-type specificity, and mechanistic integration of Tie2^+^ macrophages within TMEM doorways, these findings bridge tumor angiogenesis and metastasis research while resolving longstanding paradoxes in both fields. As TMEM doorway scores are prognostic in patients and Tie2 inhibitors are clinically tolerated, this work immediately translates to testable combination anti-metastatic strategies that could suppress breast cancer dissemination not only at its anatomic origin in the primary tumor but also at the metastatic sites where metastasis to metastasis seeding can enhance the metastatic burden ^57, 58^. TMEM doorways have been shown to be present and active at metastatic sites^57–59^, and inhibition of their opening both at primary and secondary sites could lessen overall metastatic burden and increase patient survival.

## Acknowledgments

This work was supported by the NCI grants (R01 CA255153, R01 CA240646, F32 CA243350, P30 CA013330), shared instrumentation grants (1S10OD023591-01, S10OD026852-01A1), the Gruss-Lipper Biophotonics Center, the Integrated Imaging Program, the Integrated Imaging Program for Cancer Research (IIPCR), The Evelyn Gruss-Lipper Charitable Foundation, and The Helen & Irving Spatz Family Foundation. We would like to thank the Clinical Prognostics Program in the IIPCR for autostaining support, Rotem Alon and the Analytical Imaging Facility at Albert Einstein College of Medicine for imaging support. We would also like to thank the Transgenic Mouse Core at Albert Einstein College of Medicine for help creating the Tie2-floxed mouse,

## Author contributions

Conceptualization: CLD, CRS, MHO, JSC

Methodology: CLD, CRS, XY, NDB, DC, JSC

Formal Analysis: CLD, CRS, XY, SS, DE, JSC

Software: XY, SS, DE

Investigation: CLD, CRS, PP, JH, JL, XC

Resources: DC, GSK, DE, JCM

Writing: CLD, CRS, MHO, JSC

Funding Acquisition: CLD, MHO, JSC

Supervision: DE, MHO, JSC

## Data availability statement

Data sharing is not applicable to this article as no datasets were generated or analyzed during the current study.

