## Supplemental Material for "Tie2 signaling in the tumor microenvironment orchestrates breast cancer cell dissemination through TMEM doorways"

#### Supplemental Figure Legends:

**Supplemental Figure 1: Tie2 expression in macrophages regulates VEGF-A induction by Ang2. A)** Western blot with antibodies against Tie2 and GAPDH in lysates from BAC1.2F5 macrophages, bone marrow macrophages (BMM) with low Tie2 expression (BMM Tie2 lo), and BMMs with high Tie2 expression (BMM Tie2 Hi), and HUVECs. BMMs were transduced to overexpress Tie2 (Tie2 Hi) or a control (Tie2 lo) as described in the methods section. **B)** The amount of Tie2 expression quantified by measuring the band intensity in the western blots from in (A), normalizing to GAPDH expression for each sample. n=3 western blotting experiments analyzed. ns=not significant, ***p<0.001 analyzed by one-way ANOVA and Tukey's multiple comparisons test. **C)** Immunofluorescence staining of VEGF-A in BMM Tie2 Lo and BMM Tie2 Hi macrophages, stimulated with control or Ang2 (500ng/ml) for 2 hours. Scale bars 5 µm. **D)** Quantification of immunofluorescence intensity of VEGF-A from cells in (C). n=3 experiments, 10 cells measure per experiment, ****p<0.001 analyzed by two-way ANOVA.

**Supplemental Figure 2: Tie2 knockdown in BMMs decreases VEGF-A mRNA expression and Ang2 increases VEGF-A expression, but not secretion in macrophages. A)** Western blot of Tie2 and GAPDH expression in lysates from HUVECs, Tie2 Hi BMMs treated with siRNA targeting either a control sequence (Ctrl siRNA) or Tie2 (Tie2 siRNA). **B)** Quantification of Tie2 expression levels, normalized to GAPDH, and set relative to Ctrl siRNA Tie2 expression from the western blots in (A). n=3 western blotting experiments, *p<0.05 analyzed by Student's *t*-test. **C)** Fold change VEGF-A mRNA expression from qPCR experiments. BMMs with low and high Tie2 expression (BMM Tie2 lo and BMM Tie2 hi) were pre-treated with control (DMSO) or 50 nM rebastinib for 45 minutes and then stimulated with or without 250 ng/ml Ang2 for 2 hours. n=3 individual experiments, ns=not significant, *p<0.05, ***p<0.001, ****p< 0.0001 analyzed by two-way ANOVA and Sidak's multiple comparisons test. **D)** ELISA determination of VEGF-A concentration (pg/mL) in medium conditioned by BMM Tie2 Lo and Tie2 Hi macrophages from

(C). n=3 individual experiments, performed in duplicate, **$p<0.01$ analyzed by two-way ANOVA with Sidak's multiple comparisons test. **E)** Quantification of the distance of Tie2$^+$/Iba1$^+$ macrophages from TMEM doorways in HT17 tumor sections stained in main Figure 4B, with split y-axis. Main figure 4E shows the same quantification without the split y-axis. Distances from 309 Tie2$^+$/Iba1$^+$ macrophages measured in 7 mice. Distances are binned every 10 μM, from 0 (at TMEM doorway) to 9 μM, 10 μm to 19 μM, 20 μm to 29 μm, etc.

**Supplemental Figure 3: CRISPR-Cas9 generation of the Tie2$^{fl}$ (floxed) allele in FVB mice.** **A)** Schematic of the wild-type Tie2 locus (*Tek*) on the FVB background. Exon 1 is indicated as a filled box; upstream promoter/5′ UTR and intron 1 are shown as lines. Two CRISPR guide RNAs were designed to target the upstream region (gRNA 64-41; GAAAACTTTAAGCTTGGTAT TGG) and intron 1 (gRNA IN1 72-65; CTCTCTAGAGGTGCCACTAC AGG) of Tie2. **B)** Strategy for CRISPR-Cas9-mediated insertion of loxP sites flanking exon 1. Cas9 protein, the two gRNAs, and single-stranded homology-directed repair (HDR) donors (Tie2 5′ HRD and Tie2 3′ HRD), each containing a loxP site flanked by ~40-nt homology arms, were co-injected into fertilized FVB eggs. Homology-directed repair at each cut site introduced a 5′ loxP immediately upstream of exon 1 and a 3′ loxP within intron 1, generating a floxed exon 1. Structure of the Tie2$^{fl}$ allele shown which contains loxP sites flanking exon 1 (loxP-Exon 1-loxP). Sequencing around loxP insertion sites confirms the floxing of exon 1. **C)** Diagram of Cre-mediated targeting of floxed Tie2 exon 1 (Tie2$^{fl}$) at loxP sites. **D)** PCR analysis of genomic DNA from wild-type (Tie2$^{+/+}$), heterozygous Tie2-floxed (Tie2$^{+/fl}$), and homozygous Tie2-floxed (Tie2$^{fl/fl}$) FVB mice. **E)** Bone marrow derived macrophages were isolated from Tie2$^{+/+}$ and Tie2$^{fl/fl}$ mice and cultured with 2 μm 4-OH tamoxifen for 0, 24, and 72 hours. Lysates were used for western blotting with antibodies against Tie2 and GAPDH.

**Supplemental Figure 4: Tie2 is knocked out in macrophages in the tumor microenvironment using *Csf1r-Mer-iCre-Mer* mouse model. A)** Tamoxifen treatment time course experimental model for inducible, conditional of Tie2 knockout in macrophages. PyMT tumor pieces were orthotopically transplanted into Tie2$^{fl/fl}$/*Csf1r-Mer-iCre-Mer*$^{+}$ (Cre+) and Tie2$^{fl/fl}$/*Csf1r-Mer-iCre-Mer*$^{-}$ mice and once tumors reached at least 0.6cm$^3$, mice were divided into four treatment groups. Mice were treated with corn oil (-Tam, 0hr Tam) or 50 mg/kg of 4-OH tamoxifen (+Tam) by oral gavage for 96 and 72 hours (96hr, green arrows), 72 and 48 hours (72 hr, orange arrows), or 48 and 24 hours (48 hr, red arrows) before sacrifice. Primary tumors were removed and digested for flow cytometry experiments as described in the methods section. **B)** Chart depicts the normalized counts of Tie2$^{+}$ macrophages (CD45$^{+}$/CD11b$^{+}$/F4/80$^{+}$/CD64$^{+}$/Tie2$^{+}$) in primary tumors from (A). Tie2$^{+}$ macrophage counts are shown for mice treated with corn oil (0 hr Tam, blue), or treated with tamoxifen for 96 and 72 hours (96 hr, green), 72 and 48 hours (72 hr, orange), or 48 and 24 hours (48 hr, red), showing a significant decrease in Tie2$^{+}$ macrophages as tamoxifen treatment increases. **C-J)** Graphs depict fold change in Tie2$^{+}$ C) macrophages, D) endothelial cells, E) monocytes, F) MDSCs, G) neutrophils, H) dendritic cells, and I) tumor cells compared to control mice (-Cre, -Tam) from mice treated with corn oil (-Tam) or tamoxifen (+Tam) as described in (A).

**Supplemental Figure 5: Flow cytometry panel for detection of Tie2 expression in cells of the tumor microenvironment. A)** Gating of strategy of non-myeloid (CD45$^{-}$) and myeloid (CD45$^{+}$) populations in PyMT tumors from Tie2$^{fl/fl}$/*Csf1r-Mer-iCre-Mer*$^{-}$ *and* Tie2$^{fl/fl}$/*Csf1r-Mer-iCre-Mer*$^{+}$ mice treated with corn oil or tamoxifen in Figure 6C. After gating for single and live cells (Zombie UV$^{-}$), cells were designated as tumor cells (CD45$^{-}$/EpCAM$^{+}$), endothelial cells (CD45$^{-}$/CD31$^{+}$), neutrophils (CD45$^{+}$/CD11b$^{+}$/Ly-6G$^{+}$/Ly-6C$^{-}$), monocytes (CD45$^{+}$/CD11b$^{+}$/F4/80$^{-}$/CD115$^{+}$), macrophages (CD45$^{+}$/CD11b$^{+}$/F4/80$^{+}$/CD64$^{+}$/Ly-6G$^{-}$), dendritic cells (CD45$^{+}$/CD11c$^{+}$/Ly-6C$^{-}$/F4/80$^{-}$/CD64$^{-}$), and myeloid derived suppressor cells (MDSCs; (CD45$^{+}$/CD11b$^{+}$/Ly-6G$^{-}$/CD115$^{+}$). **B)** PE-IgG, in place of Tie2-PE-IgG antibody, and a Tie2 FMO (fluorescence minus one) were used to gate for Tie2$^{+}$ cells, as they show no positive Tie2 signal in the gate chosen, compared to $Cre^+$ control mouse (panels on right). **C)** $Tie2^+$ macrophage counts from parental macrophage population ($CD45^+CD11b^+F4/80^+CD64^+$) in mice treated with corn oil and stained with Tie2 antibodies ($Tie2^+$ cells, pink population) or stained with Tie2 isotype control antibodes (Isotype Ctrl, gray population). Dashed lined indicates where the gate for $Tie2^+$ and $Tie2^-$ signal was placed based on the isotype control staining.

**Supplemental Figure 6: Average number of F4/80 and Ly-6G expressing cells in PyMT/Tie2$^{fl/fl}$/*Csf1r-Mer-iCre-Mer* mouse model. A, C)** Tumor tissue sections from mice treated with tamoxifen (+Tam) or corn oil (-Tam) to knockdown Tie2 in macrophages in Figure 6C were stained for DAPI and (A) F4/80 to identify macrophages or (C) Ly/6G to identify neutrophils. **B)** Quantification of the average number of $F4/80^+$ macrophages per field from staining in (A). ns=not significant, analyzed by Student's *t*-test, n=7-8 mice per group, each dot represents the average value for a mouse. **D)** Quantification of the average number of Ly-6G$^+$ neutrophils per field from staining in (C). ns=not significant, analyzed by Student's *t*-test, n=7-8 mice per group, each dot represents the average value for a mouse.

**Supplemental Figure 7: Rebastinib and paclitaxel enhance overall survival in mice with metastatic breast cancer. A)** Diagram of short-term neo-adjuvant treatment study design. Tumor chunks from MMTV-PyMT mice were orthotopically transplanted into FVB mice. Tumors were allowed to grow until at least 0.25 $cm^3$ in volume and mice were then divided into four treatment groups: control (CreL injected *i.v.* every 5 days and Control chow), paclitaxel (10 mg/mL *i.v.* every five days), rebastinib chow (22 mg/kg *ad libitum*), and paclitaxel and rebastinib chow. Mouse were treated continuously until tumors from control treated mice reached 2 $cm^3$ at which point all mice were sacrificed and CTCs were collected. **B)** Fold change quantification of circulating tumor cells from mice in (D). *$p<0.05$, **$p<0.01$, analyzed by a one-way ANOVA, n=4-5 mice per group, each dot represents the normalized value for a mouse.

**Supplemental Figure 1: Tie2 expression in macrophages correlates with VEGF-A induction by Ang2**

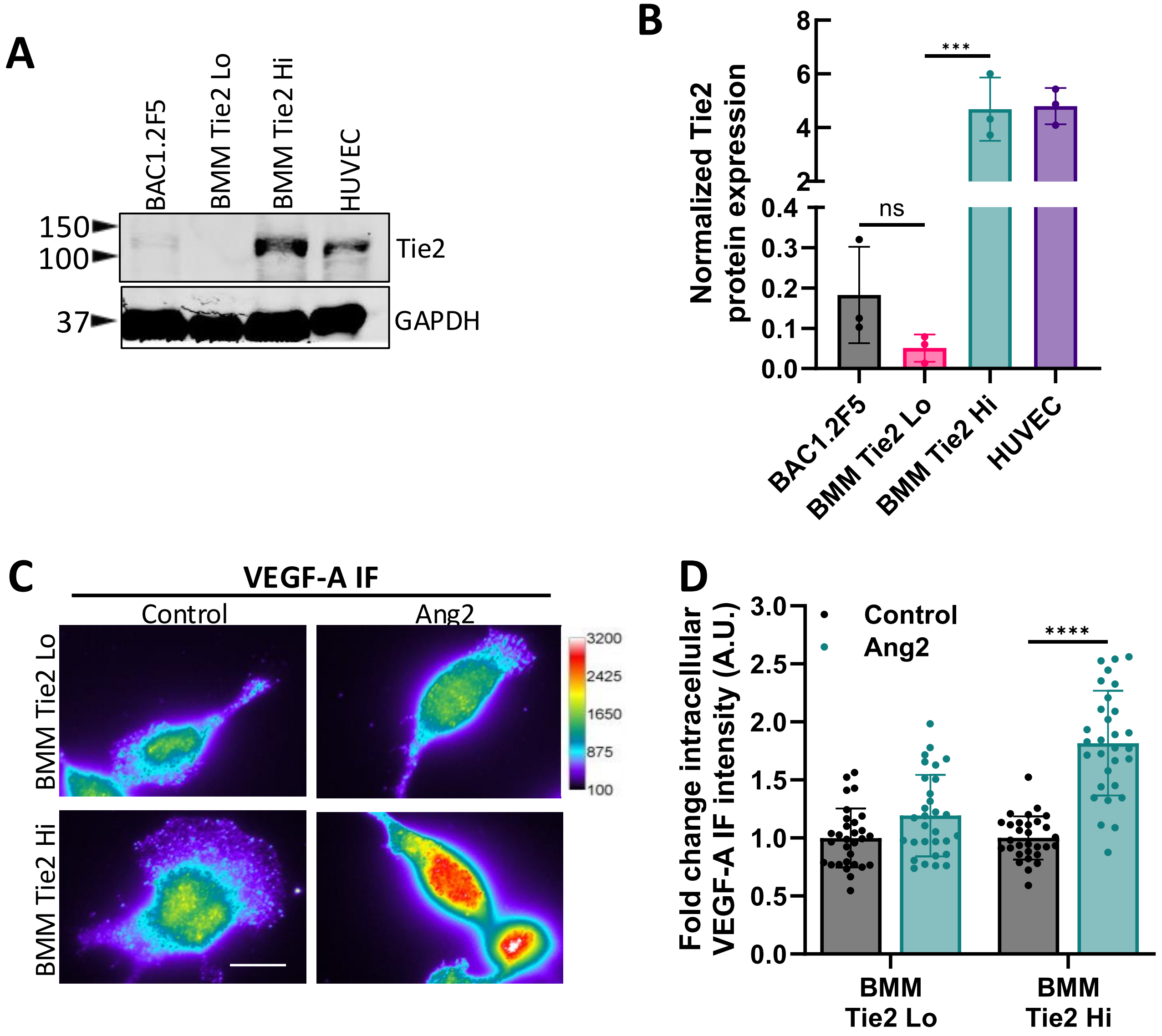

**Supplemental Figure 2: Tie2 knockdown in BMMs decreases VEGF-A mRNA expression**

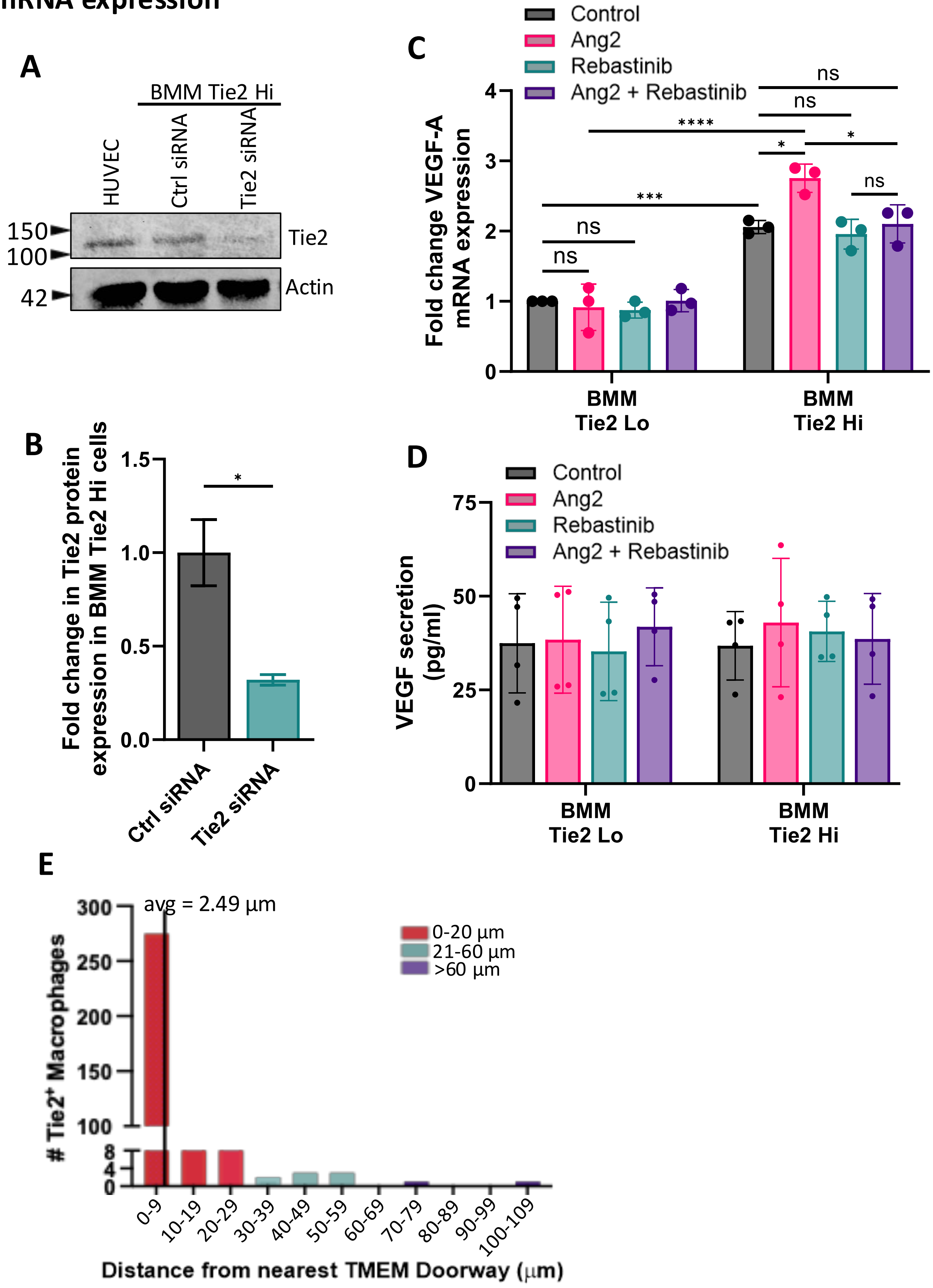

#### Supplemental Figure 3: CRISPR/Cas9 generation of the Tie2$^{fl/fl}$ allele in FVB mouse

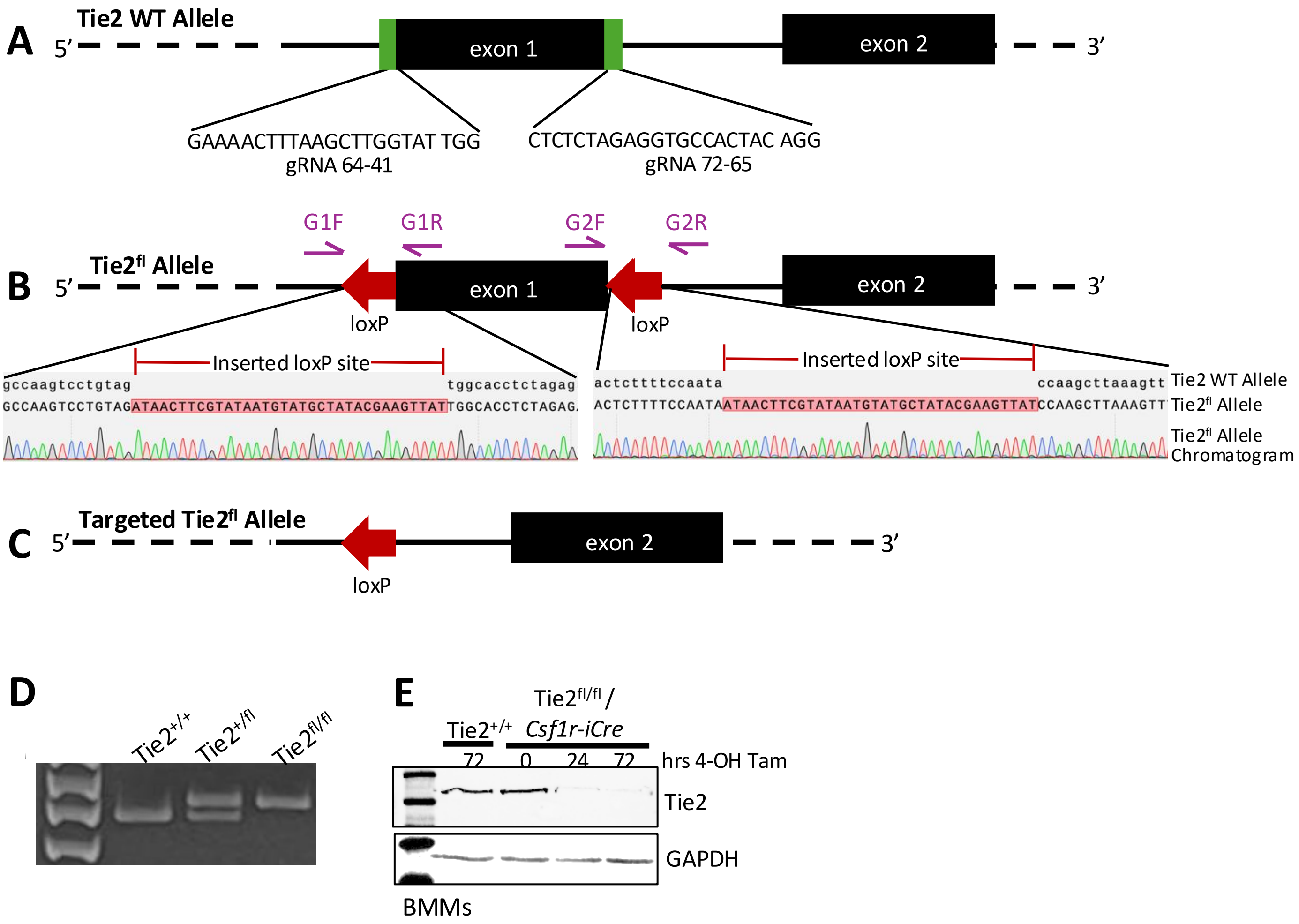

### Supplemental Figure 4: Tie2 is knocked out in macrophages in the tumor microenvironment using *Csf1r-Mer-iCre-Mer* mouse model

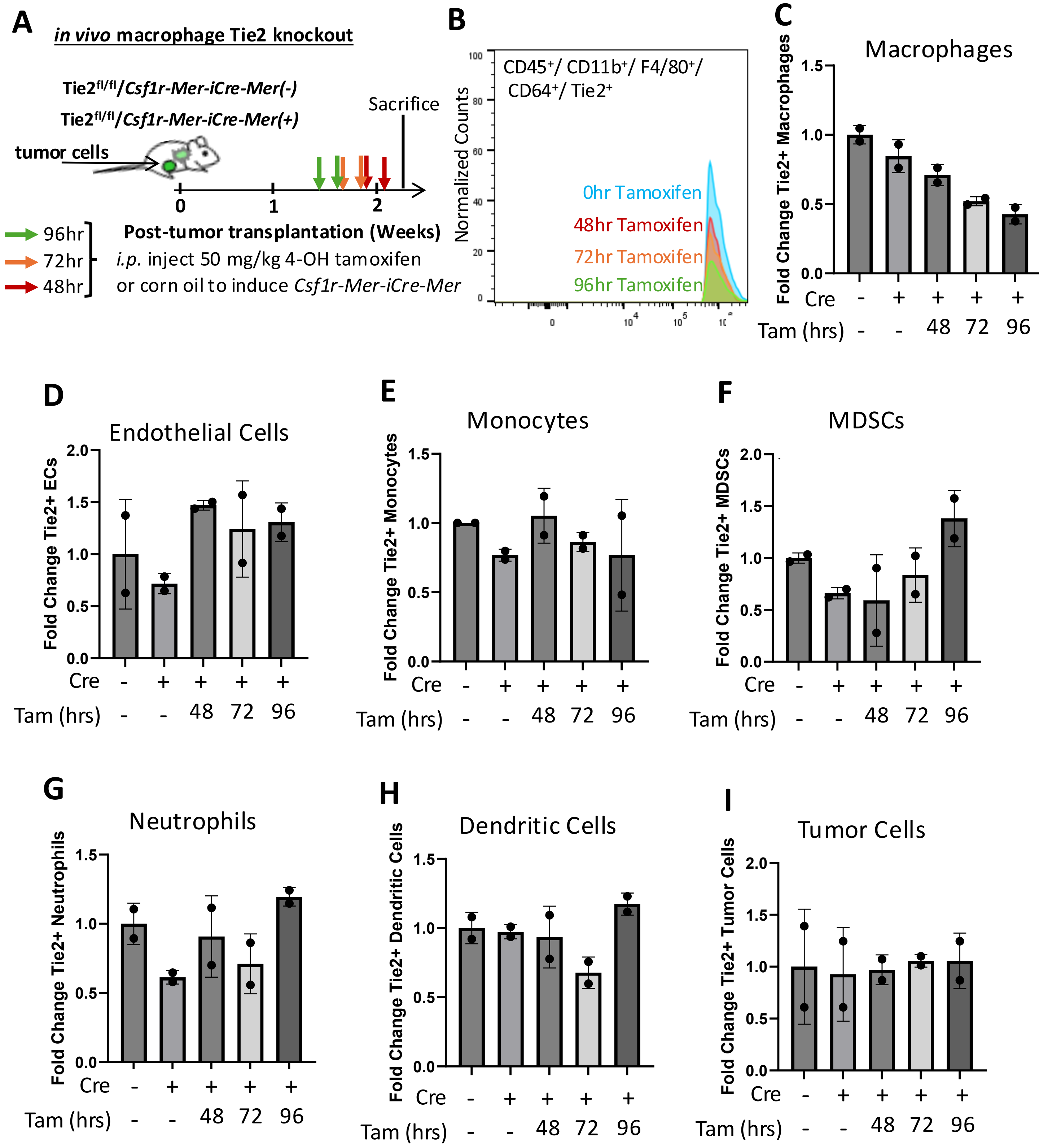

**Supplemental Figure 5: Flow cytometry panel for detection of Tie2 expression in cells of the tumor microenvironment**

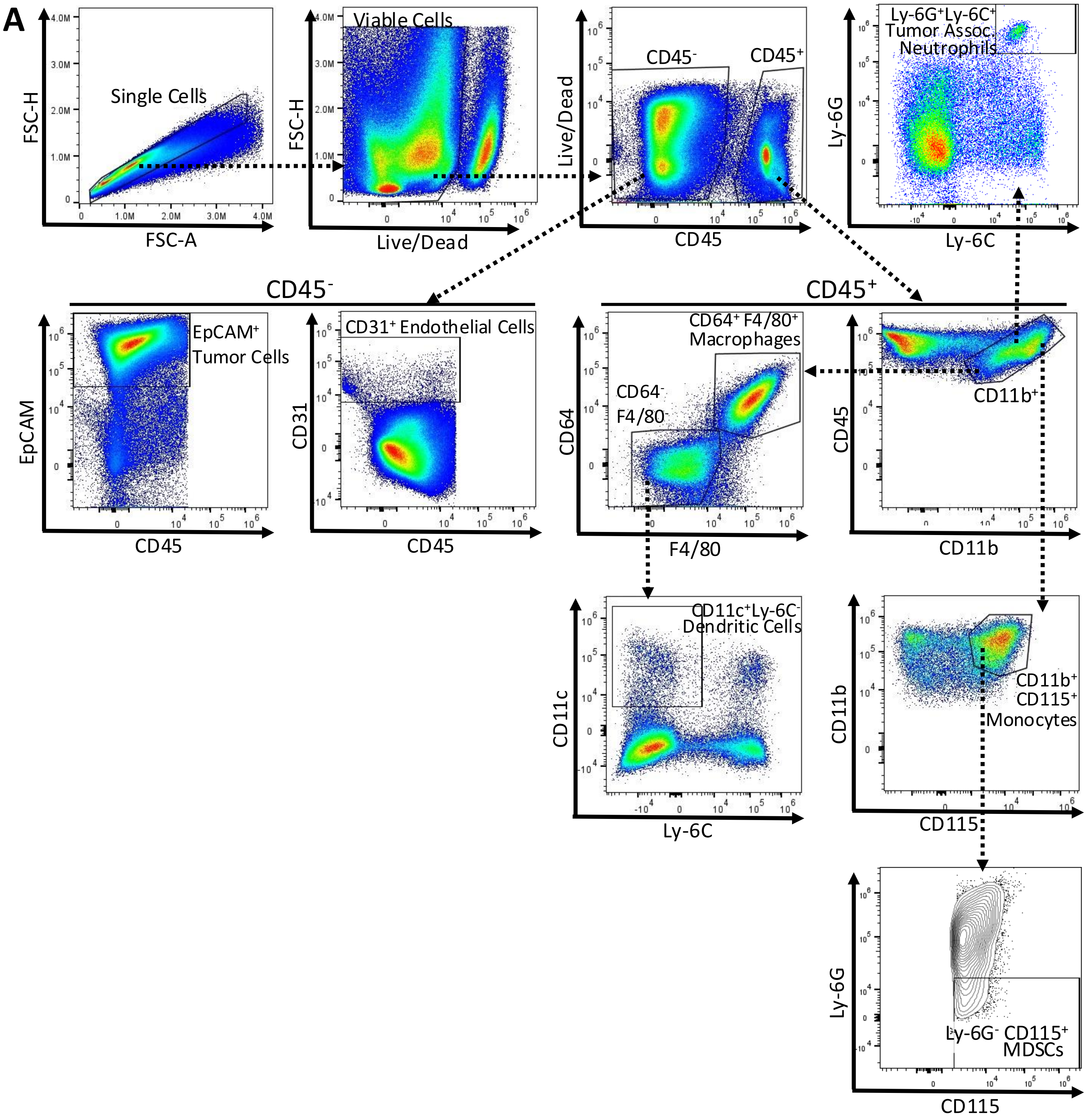

### Supplemental Figure 5: Flow cytometry gating for Tie2 expression in macrophages in the tumor lysates

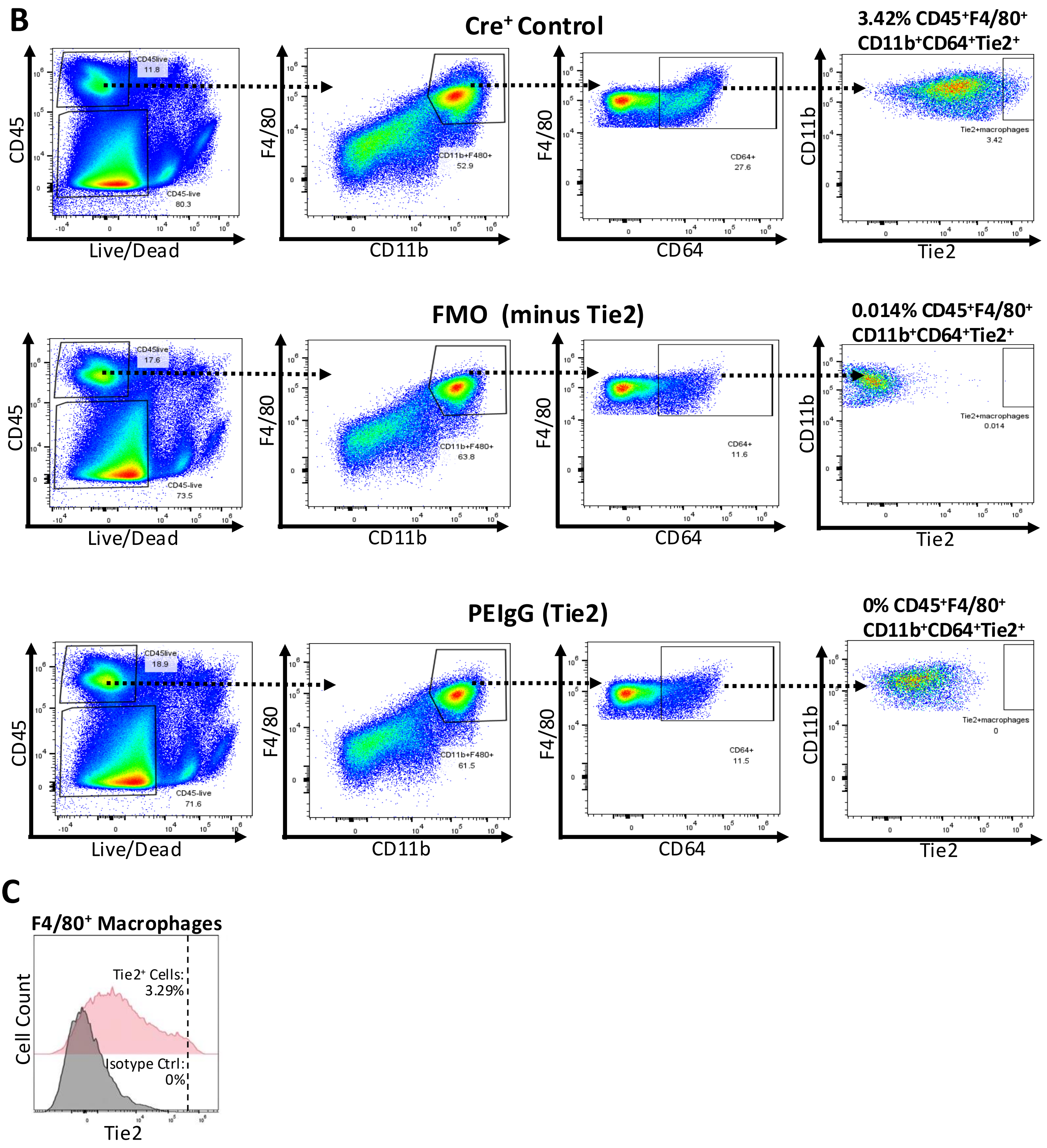

**Supplemental Figure 6: Average numbers of macrophages and neutrophils in PyMT/Tie2$^{fl/fl}$/*Csf1r-Mer-iCre-Mer* model**

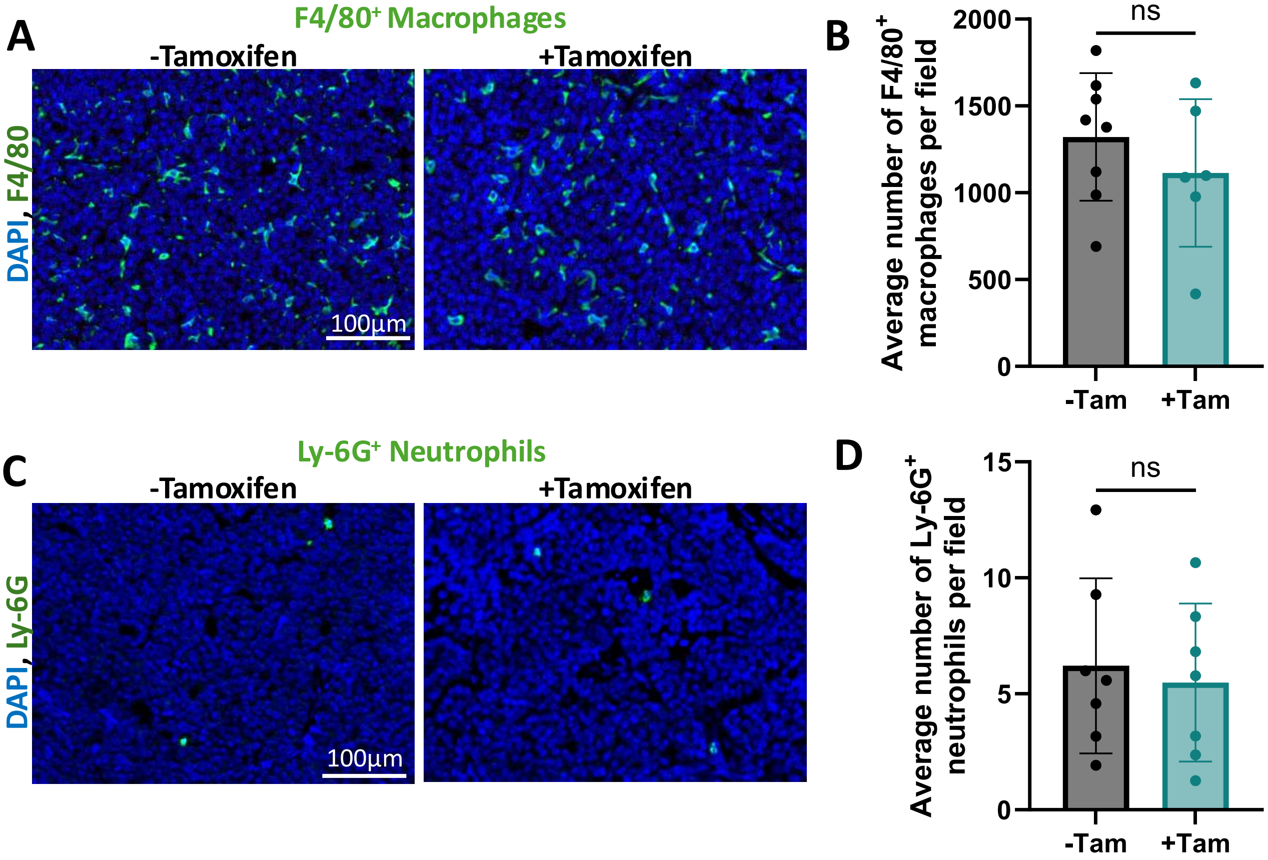

**Supplemental Figure 7: Rebastinib inhibits paclitaxel-induced influx of CTCs**

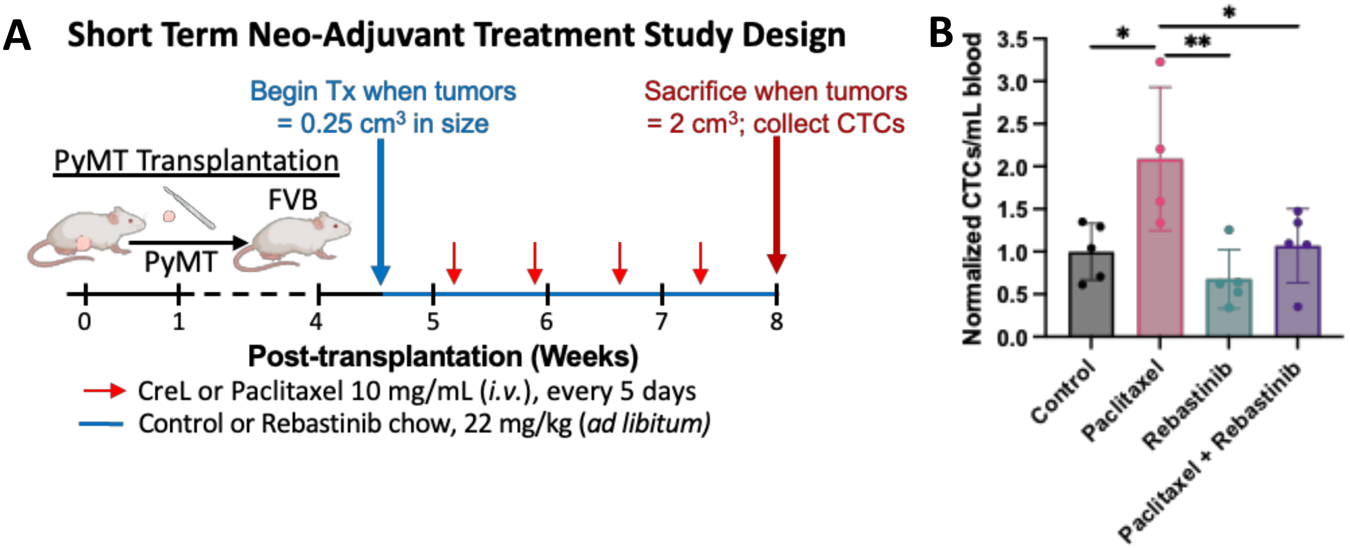
